# Predicting Protein Function and Orientation on a Gold Nanoparticle Surface Using a Residue-Based Affinity Scale

**DOI:** 10.1101/2022.03.29.486298

**Authors:** Joanna Xiuzhu Xu, Md Siddik Alom, Rahul Yadav, Nicholas C. Fitzkee

## Abstract

The orientation adopted by proteins on nanoparticle surfaces determines the nanoparticle’s bioactivity and its interactions with living systems. Here, we present a residue-based affinity scale for predicting protein orientation on citrate-gold nanoparticles (AuNPs). Competitive binding between protein variants accounts for thermodynamic and kinetic aspects of adsorption in this scale. For hydrophobic residues, the steric considerations dominate, whereas electrostatic interactions are critical for hydrophilic residues. The scale rationalizes the well-defined binding orientation of the small GB3 protein, and it subsequently predicts the orientation and active site accessibility of two enzymes on AuNPs. Additionally, our approach accounts for the AuNP-bound activity of five out of six additional enzymes from the literature. The model developed here enables high-throughput predictions of protein behavior on nanoparticles, and it enhances our understanding of protein orientation in the biomolecular corona, which should greatly enhance the performance and safety of nanomedicines used in vivo.

---

Understanding the behavior of surface-adsorbed proteins on nanoparticles (NP) is crucial when NPs are used in vivo as a therapeutic drug carrier, phototherapeutic agent, or imaging tool^1–4^. There is increasing evidence that the composition of the adsorbed proteins is not as important as their structure and orientation^5–7^, as the latter governs the three-dimensional presentation of proteins for molecular recognition, which will define the biological performance and fate of the NP^8–10^. For example, the arrangement and orientation of the adsorbed proteins on a NP define the distribution of epitopes presented for immune response^5^ and determine the success of cell targeting^6,9^. Indeed, steric hindrance from the NP surface can restrict access to the important sites of the adsorbed protein, preventing its molecular recognition^11^. In this regard, various methods of controlling the bound protein orientation have been developed for retaining protein bioactivity and enhancing targeting efficiency of protein-NP conjugates^10,12–14^. While knowing the bound orientation is critical to predicting the fate of the NP, obtaining this information experimentally is a daunting task. Only a handful of NMR techniques can probe the NP-binding sites of a protein in situ^15,16^, but these methods have substantial technical challenges and limitations^17^.

In this work, we hypothesize that protein-NP association is mediated primarily by the surface residues of a protein. Therefore, we generate an experimental affinity scale for each residue by varying a key binding position in the small GB3 protein^18^. Competitive protein binding is required to rationalize residue differences^19,20^, demonstrating that both kinetic and thermodynamic considerations influence adsorption^21–23^. This affinity scale and a simple surface-area based algorithm predict the preferred orientation/function for three proteins presented here, including two blind predictions; moreover, our findings rationalize the observed function of five additional proteins from the literature. The presented approach is widely applicable to different nanoparticle systems to improve performance and biosafety, and it enables high-throughput predictions in large protein datasets.

## A residue-based affinity scale for AuNP binding

We began by systematically examining a series of GB3 variants where the 13^th^ position was changed to all 20 amino acids (**Fig. 1a**). Prior work established that the interaction of GB3 with 15-nm citrate-capped AuNP is mediated by lysine residues^18,24^. In particular, K13 contributes most to the binding, and mutation of this residue to alanine significantly reduces GB3 binding^18^. None of the variants had significantly altered structure or stability as assessed by NMR (**Supplementary Information**). To measure the binding affinity of residue X in regard to the simplest residue glycine (G), an equal amount of K13X GB3 was mixed with K13G GB3 before exposing to AuNPs for competitive binding (**Fig. 1b,c**). After equilibrium is attained (1h), the reduction in NMR signals corresponds to the amount of GB3 in the hard corona, as all signals from the bound proteins are broadened beyond detection (**Fig. 1c, right**)^19^. To disentangle NMR signals of the protein mixture, variant K13X GB3 is ^15^N-labeled, whereas the reference variant K13G GB3 is ^13^C-labeled. This enables in situ quantification of the bound amount of each variant, and the affinity scale for all 20 amino acid residues. The *α* value is defined as the ratio of the K13X variant bound relative to K13G (**Table 1**). Thus, *α* is derived using non-disruptive, in situ measurements, where the corona is formed in the presence of protein competition.

**Table 1.**
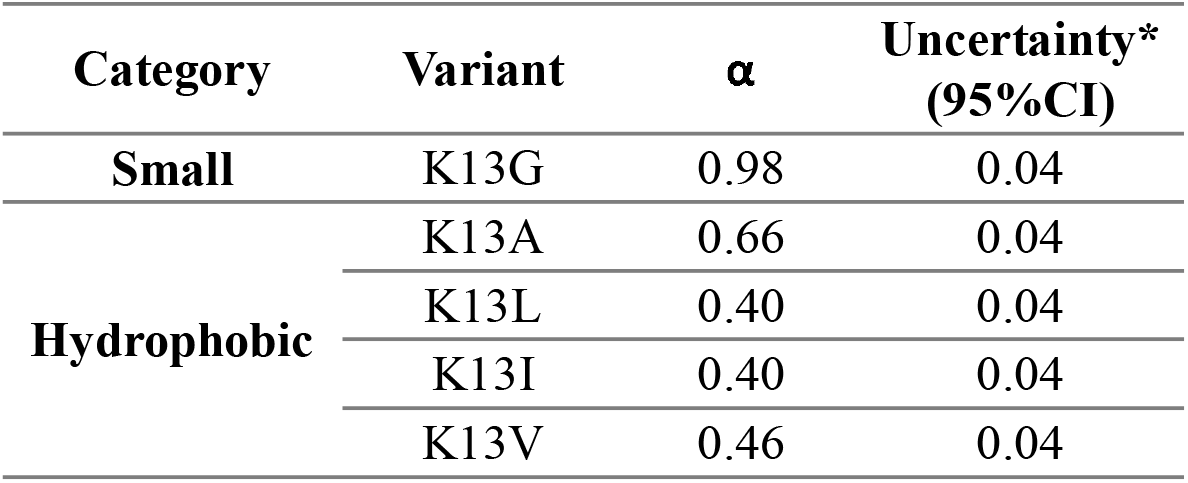
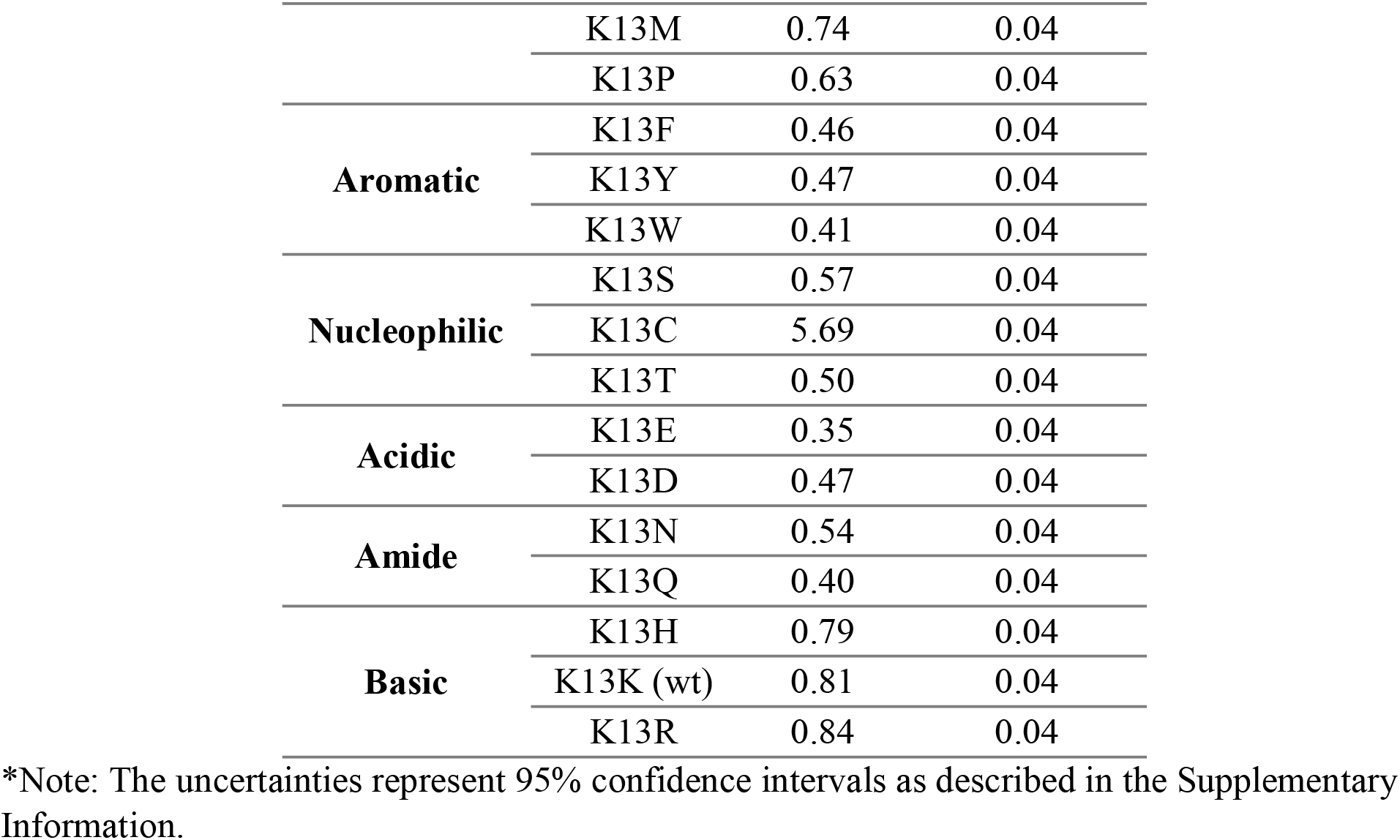
Alpha values of 20 GB3 variants

**Fig. 1.**
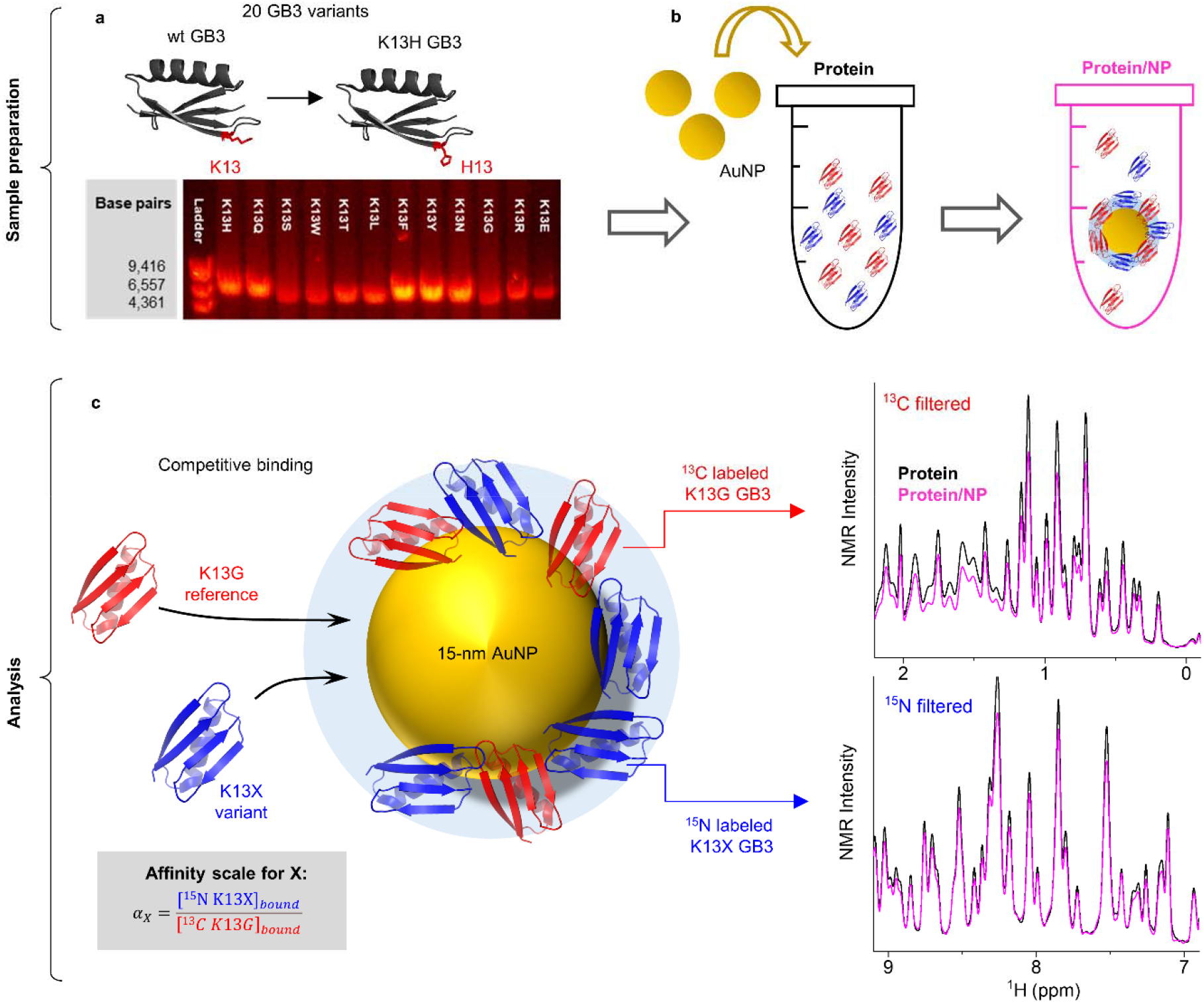
Determination of affinity scales for 20 amino acid residues with host-guest competitive binding experiments. **a**, Mutagenesis of K13 into each of the 20 amino acids with histidine (H13) as an example. The agarose gel electrophoresis shows several examples of PCR products for GB3 mutants. **b**, Preparation of competitive binding experiments. Two GB3 variants (protein) are mixed before adding AuNPs (protein/NP sample), and a protein corona is formed upon adsorption. **c**, NMR analysis for quantification of binding capacity of each variant. The reference variant ^13^C labeled K13G and ^15^N labeled K13X is measured with ^13^C filtered and ^15^N filtered 1D NMR, respectively. The bound concentration of each variant is deduced from NMR signal reduction from the protein sample (black) and protein/NP sample (magenta) prepared in **b**. Finally, the affinity scale (*α*_*X*_) for residue X is the ratio between the bound concentrations of K13X ([^15^*N* K13X]_*bound*_) and K13G ([^13^*C* K13G]_*bound*_)

The fact that *α*_*G*_ (^15^N K13G vs. ^13^C K13G) equals 1 (*α*_*G*_ = 0.98 ±0.04) indicates different isotopic labeling has no measurable impact on the GB3-AuNP interaction. Cysteine has the highest affinity towards AuNP (*α*_*C*_=5.69±0.04), due to strong Au-S bond formation^25^. All other α values are smaller than 1, regardless of electrostatic, geometric, or hydrophobic properties of a residue. This is surprising, and it suggests that reducing steric bulk and maximizing protein-surface contact can be advantageous to adsorption. The fact that the smallest side chain renders K13G GB3 highest affinity in competition led us to evaluate the role of the steric effect afforded by different residues.

Indeed, the *α* values correlate inversely with amino acid molecular volumes (AA volume) among K13X GB3 variants modified with aliphatic side chains (**Fig. 2a**) and hydrogen bonding (H bond)/aromatic side chains (**Fig. 2b**), with strong linearity observed for each residue sub-group (**Fig. 2d** and **2e**), except K13Q. This steric effect appears most significant for smaller amino acid side chains. The slopes from the aliphatic subgroup and the polar/aromatic subgroup differ by a factor of five ((−49 ± 7) × 10^−4^ vs. (−9.6 ± 2) × 10^−4^, **Fig. 2d,e**). Besides this, the hydrophilicity afforded by the terminal hydroxyl group (S, T, Y), or the polarity (W) may play a role in counteracting the increasing steric effect in the polar/aromatic subgroup. Electrostatic interactions can also counter the steric effect, as demonstrated in the charged subgroup (**Fig. 2c**).

**Fig. 2.**
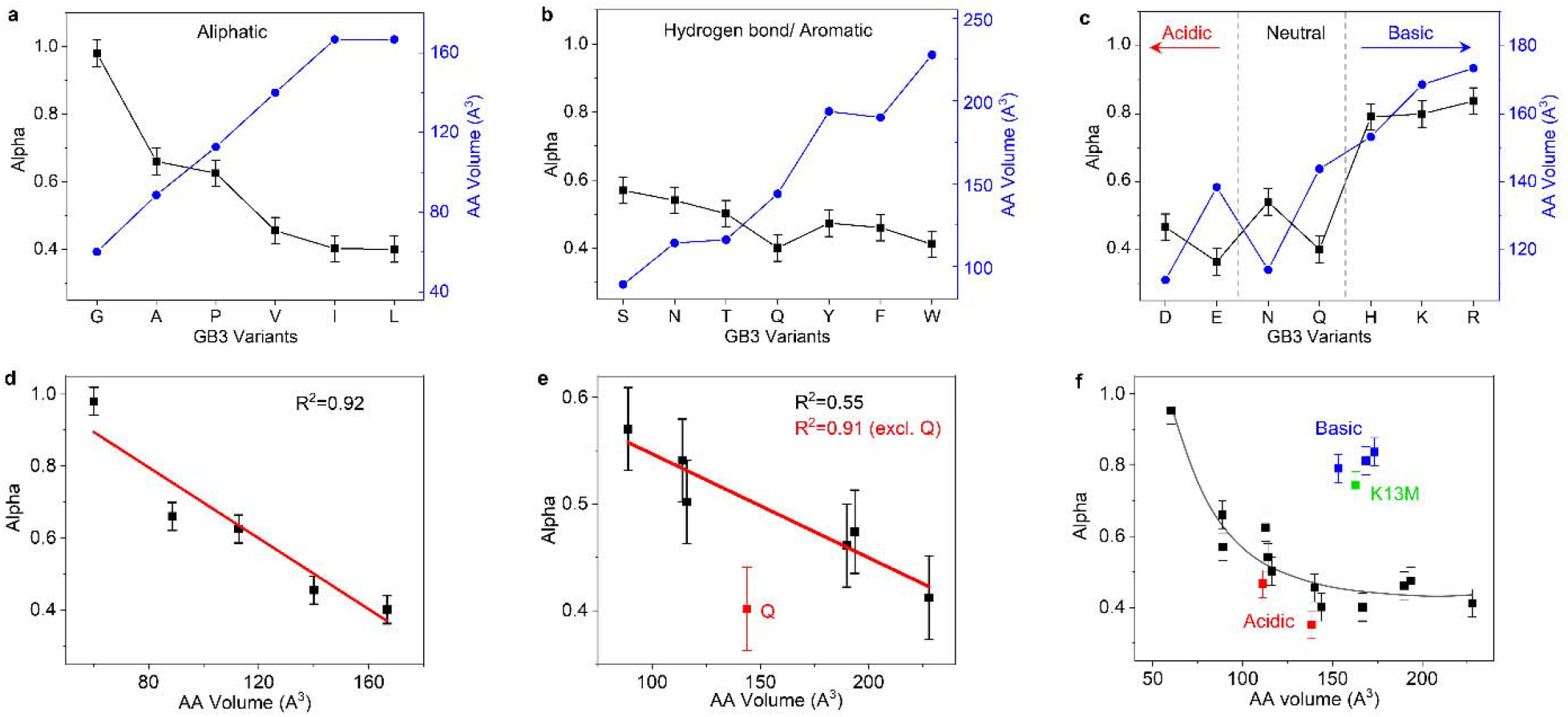
Effect of residue identity on binding affinity. Correlation of *α* value (black) and amino acid (AA) molecular volumes^26^ (blue) for K13X variants with aliphatic side chains (**a**), hydrogen-binding/aromatic side chains (**b**), and electrostatic side chains (**c**). The red arrow and blue arrow indicate increasing acidity and basicity. **d, e** Linear fitting of alpha values as a function of AA volumes for variants presented in **a** and **b. f**, Relationship of alpha values and AA volume for all 20 GB3 variants. The grey curve is a polynomial fit to guide the eye. Data points of acidic, basic residues and K13M are highlighted in red, blue and green, respectively. All error bars are an averaged 95% confidence interval error as illustrated in the **Supplementary Information**.

While H, K, and R have a ~30% larger volume than D or E, their alpha value is much larger, reflecting a favorable interaction between these residues and the negatively charged citrate-coated surface. This effect is clearly seen in histidine, where increasing pH leads to a steady decrease in *α*_*H*_ (**Supplementary Information, Fig. 6**). However, even for D and E, an increase in volume correlates with a decrease in alpha (**Fig. 2c**), and this trend applies for residues with similar functional groups (e.g. Q vs. N). An unexpectedly high *α* is observed for K13M, which may be due to strong attraction between organic sulfides and AuNPs^27^. Overall, increasing side chain volumes appears to reduce competitive binding in our host-guest system, but this effect diminishes for larger side chains (**Fig. 2f**). Larger side chains on a protein’s surface can sample more rotomeric states, decreasing the protein’s ability to maximize contact with a NP surface, and this may explain the trend we observe.

## Competition is essential for predicting binding affinity

Next, we investigated the thermodynamic and kinetic aspects of adsorption as they relate to residue alpha values. First, a single GB3 variant was titrated into an AuNP solution to quantify thermodynamic dissociation constants (*K*_*d*_) of five selected GB3 variants, which represent distinct properties and a range of alpha values: K13G – smallest; K13A – aliphatic; K13K (WT) – basic; K13E – acidic; and K13Y – aromatic. The increasing red shift of the AuNP plasmonic peak is caused by increasing protein adsorption; this shift can be fit with the Langmuir isotherm (**Fig. 3a,b**). Surprisingly, there is no statistical difference between thermodynamic *K*_*d*_ values for these variants. This is in stark contrast to their different affinity scales (**Fig. 3c**). Subsequently, we investigated adsorption kinetics of these variants using time-resolved SOFAST-HMQC 2D NMR spectroscopy, where signal reduction corresponds to protein binding. Again, no difference was observed for binding kinetics or capacity among these variants when only a single variant was present, e.g. K13G and K13E (**Fig. 3e** and **Supplementary Fig. 9**). However, when competition is introduced by exposing a mixture of two variants to a solution of AuNPs, K13G was adsorbed faster than K13E and bound to a larger degree, corresponding to its lower alpha value. Note that only a few well-resolved peaks (e.g., G9, T11, **Fig. 3d** and **Supplementary Fig. 7**) can be used for quantification of binding in this mixture (**Fig. 3f**) due to significant signal overlap^19^, and this approach is somewhat less precise than the ^13^C, ^15^N differential labeling method used above. In these competition experiments, the total protein bound remains the same as when individual variants are bound, supporting the alpha value approach (**Fig. 3e** vs. **3f**). Together, these results emphasize the importance of competition. In vivo, proteins simultaneously compete for a NP surface, and multiple thermodynamic and kinetic aspects are involved;^21–23,28^ the alpha value captures these complexities in one simple metric. Our results also address the limitations of an individual protein binding approach, which may not accurately reflect the corona in complex protein mixtures^7,29,30^.

**Fig. 3.**
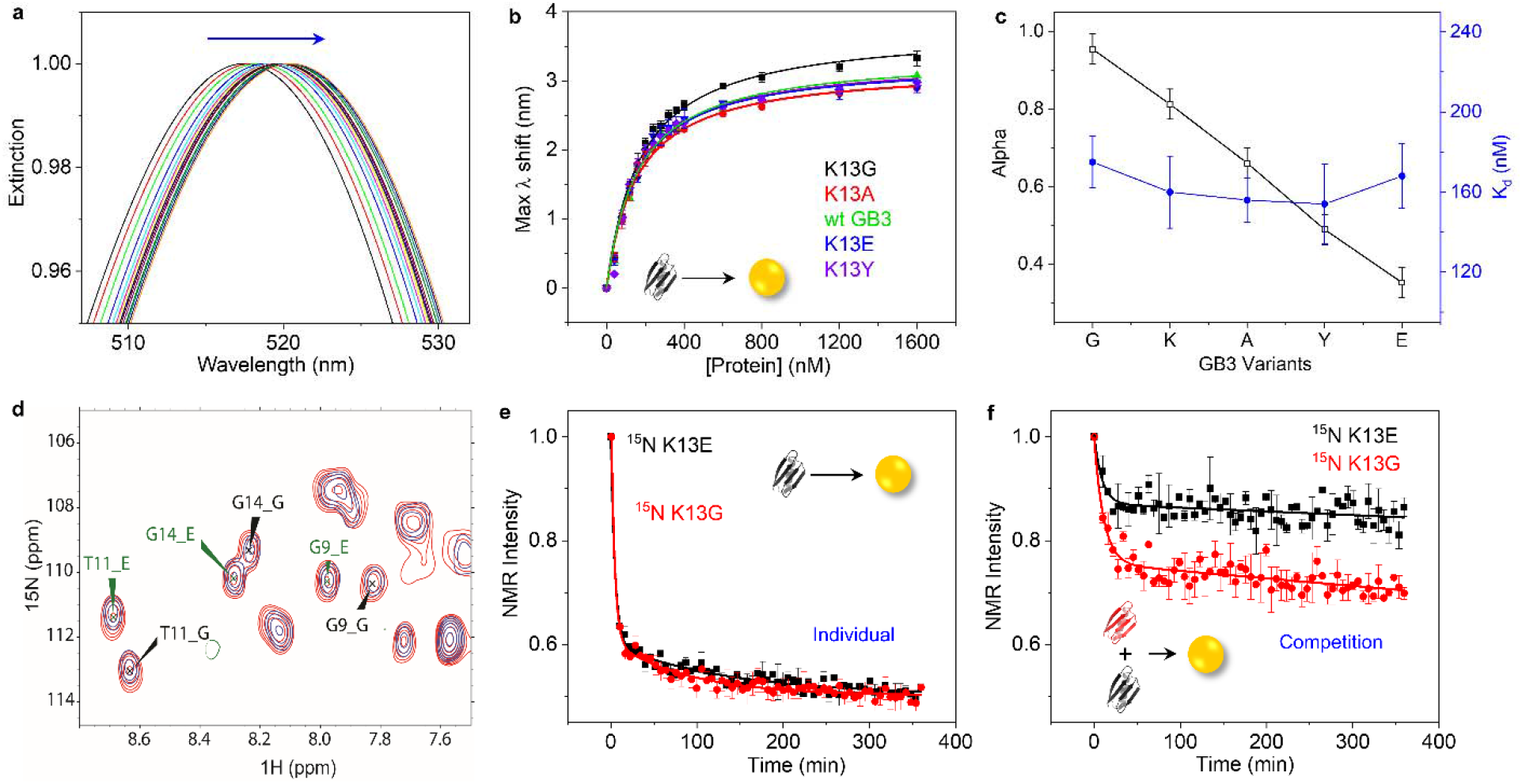
Role of competition in protein corona formation. **a**, UV-vis titrations of an individual variant into AuNPs, using K13Y GB3 as an example. The arrow represents increasing red shifts of the AuNP plasmonic peak caused by enhanced K13Y GB3 binding. **b**, Maximum wavelength shifts of the AuNP plasmonic peak in **a** plotted against protein concentrations for five representative GB3 variants. Each data set is fit with Langmuir adsorption isotherm model. The error bars represent deviation from two replicates. **c**, A comparison of alpha values obtained from competitive binding versus Langmuir dissociation constants (*K*_*d*_) obtained from **b**. The error bars for *K*_*d*_ are the fitting errors. **d**, Example SOFAST-HMQC NMR spectrum for kinetic binding study of two GB3 variants binding onto AuNP in competition. The peaks labeled in black (and ending in “_G”) are from K13G and those in green (and ending in “_E”) are from K13E GB3. The red spectrum represents a protein control sample without AuNPs (0 min point), and the blue spectra shows the protein/AuNP sample after binding equilibrium is reached (365 min). Peak intensities are reduced due to binding. Complete spectra are presented in **Supplementary Fig. 7. e**, Protein peak intensity change as a function of incubation time with AuNPs for K13E (black) and K13G (red) GB3, respectively. Upon mixing an individual protein with AuNPs, a SOFAST-HMQC spectrum was collected every 5 mins up to 365 mins. The average peak intensity at each time point is normalized by that of the protein control sample and calibrated with the external reference^19^. An example SOFAST-HMQC NMR spectrum for individual K13E GB3 binding is shown in **Supplementary Fig. 8**, and individual binding kinetic profiles of other selected variants are showed in **Supplementary Fig. 9**. All fitting parameters are shown in **Supplementary Table 5. f**, Competitive binding kinetics of K13G versus K13E GB3, with example SOFAST-HMQC spectra presented in **d**. An equal amount of each variant is mixed before addition of AuNPs, rendering total protein concentration double that of the individual binding experiments in **e**.

## Predicting protein bound orientation and function on AuNPs

Finally, we attempted to predict the preferred AuNP-binding surface of several proteins using the alpha values. We devise a simple metric for the binding affinity of a residue, calculated by multiplying the alpha value with the accessible surface area relative to a fully exposed sidechain^31^ (**Fig. 4a**). To visualize regions of high binding affinity, a protein is overlaid on a grid, and the average binding affinity from nearby residues is calculated at each surface-accessible grid point. Grid points are represented by virtual atoms, colored by low (white) to high (red) binding affinity, and values below a threshold are not shown (full details are provided in **Supplementary Information**, and scripts are available to download on GitHub). The result for GB3 (**Fig. 4a, right**) reveals three high-affinity regions on either side of the β-sheet; conversely, the alpha helix has relatively low affinity. Based on this, we predict that GB3 interacts with AuNPs by its β-sheet face, and this orientation is consistent with previous work^18^.

**Fig. 4.**
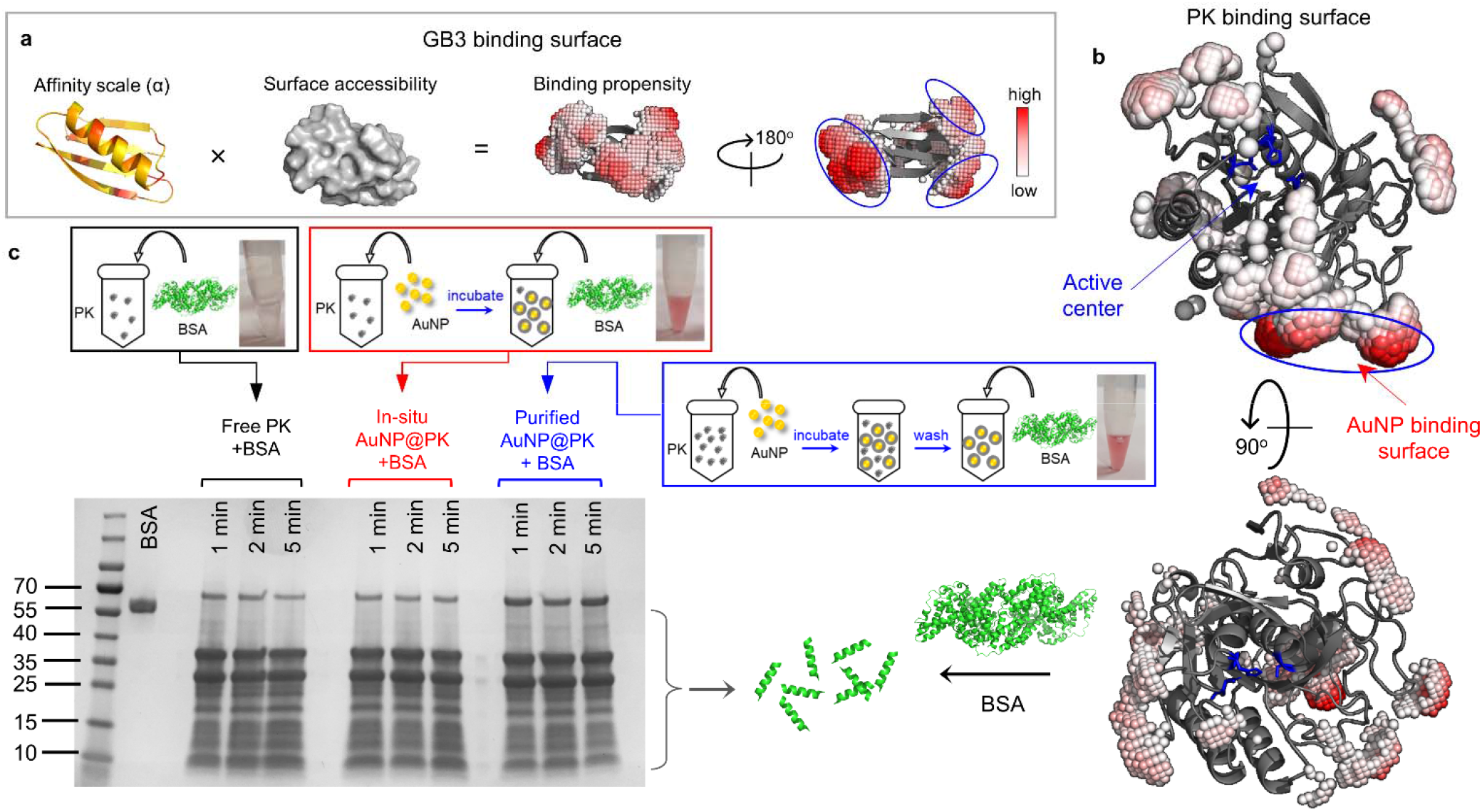
Prediction of bound orientation of Proteinase K (PK) on AuNP. **a**, A scheme for deducing the preferred bound surface of GB3 by taking both the alpha value and surface accessibility into account (see main text). Alpha values in the leftmost image are depicted using a yellow (low) to red (high) gradient. The α-helix side and β-sheet side of GB3 show the predicted AuNP-binding surface using color-coded virtual atoms, colored white (low affinity) to red (high affinity). No virtual atoms are drawn below a fixed threshold value. **b**, Cartoon structure of PK showing the predicted AuNP-binding surface circled in blue, and its active center constructed by S224, H69 and D39 as highlighted in blue sticks. **c**, Preparation and results of PK activity assay for free and AuNP-bound PK. The PK concentration in all assays is kept at 0.01 mg/mL and substrate BSA concentration is at 2 mg/mL for all samples to achieve limited proteolysis. As shown in the scheme (top), before mixing with BSA, the in-situ bound PK sample (in-situ AuNP@PK) is prepared by adding AuNPs to bind ~90% of free PK without further purification, whereas the purified AuNP@PK is washed after incubating excess amount of PK with AuNPs (**Supplementary Information**). SDS-PAGE shows limited proteolysis of BSA using free PK (lane 4-6), in-situ AuNP@PK (lane 8-10), and pre-coated and washed AuNP@PK (lane 12-14) for reaction times of 1 min, 2 mins, and 5 mins (from left to right), respectively. Lane 2 shows a 0.2 mg/mL BSA control with no PK. The associated schemes outline their sample preparations, and the photographs show the assay solutions.

Subsequently, we tested our approach for predicting the orientation of unknown proteins. We chose two enzymes for this purpose, and we hypothesized that enzyme activity could be used to monitor protein orientation indirectly: assuming that a protein remains folded and packed on the NP surface^24^, a bound orientation with a sterically hindered active site should exhibit reduced activity^6,13^. Conversely, an orientation that exposes the active site should have near-normal activity. Proteinase K (PK) has a catalytic triad constructed by S224, H69 and D39, and it non-specifically cleaves peptide bonds in a protein (active site highlighted in blue, **Fig. 4b**). Our model predicts that PK should bind to AuNPs through the residues highlighted by the red binding patches (**Fig. 4b**), leaving its substrate access sterically unaffected. We examine this prediction by quantifying the proteolytic efficiency of NP-bound PK for digesting bovine serum albumin (BSA). With a size of 66 kDa, BSA should be sensitive to any steric hindrance toward the substrate-binding site of PK. For rigorous validation, the AuNP-bound PK (AuNP@PK) was prepared in two ways: in-situ, where AuNPs are simply mixed with PK without further perturbation; and washing, where AuNP@PK was centrifuged and resuspended three times in buffer to ensure removal of free PK from the solution (purified AuNP@PK). The purified AuNP@PK was characterized by UV-vis spectrometry and DLS (**Supplementary Fig. 10**) to ensure no AuNP aggregation. The activities of both in-situ and purified AuNP@PK were compared with a control sample of PK with no AuNPs, and the total PK concentration was held constant (**Supplementary Information**). In both cases, the nanoparticles are fully saturated with PK before mixing with BSA; otherwise, BSA may bind and be protected from proteolysis^32,33^. This protection was observed when AuNPs in large excess (~ 4 times more AuNP surface) were added to conjugate a limited amount of PK, followed by mixing with BSA (**Supplementary Fig. 11**).

After optimizing the enzyme/substrate ratio for limited proteolysis of BSA within a 1-5 minute observation time, we found that PK exhibits identical proteolytic efficiency in both free and AuNP-bound (AuNP@PK) forms (**Fig. 4c**). In particular, the band intensities corresponding to the cleavage of intact BSA into smaller peptides are almost identical. Adding AuNPs to PK in the in-situ experiment shows no reduced activity, and washing the AuNP@PK particles before mixing with BSA confirms that the activity originates from the on-AuNP PK. This suggests that the orientation of PK on the AuNP surface matches our prediction, with no steric hindrance to its active site (**Fig. 4b**).

Using this approach, we predict that the AuNP binding surface and substrate binding site will be coincident in human carbonic anhydrase II (HCA). As depicted in **Fig. 5a**, the HCA active center is constructed by Zn^2+^ (gray sphere), H96, H94, and H119 (highlighted in sticks)^34^. During the conversion of carbon dioxide into carbonic acid by HCA, residue H64 (showed as sphere) plays an essential role in shuttling protons from the active center to solvent^35^. The binding of HCA onto AuNP is predicted to occlude the active center and prevent proton transportation by H64 to the environment. We tested the activity of HCA in the free and bound states using a *p*-nitrophenyl acetate (*p*NPA) hydrolysis assay (**Fig. 5b**), where NP-bound HCA (AuNP@HCA) was prepared in-situ and by AuNP washing, similarly to PK. Free HCA converts *p*NPA to *p*NP rapidly and efficiently upon mixing, as indicated by the increasing absorbance from *p*NP at 404 nm (**Fig. 5c**). However, on AuNPs, the activity of the bound HCA is mostly inhibited regardless of preparation method (**Fig. 5d and 5e**). Prior work by Saada *et al*. suggests that this inactivation occurs mainly due reduced accessibility of H64 rather than denaturation of HCA by AuNP^36^. In their work, the authors show that introducing functional groups with histidine-like imidazole rings on AuNP could support the activity of the adsorbed HCA. This result also suggests the channel of proton shuttling is oriented toward the AuNP surface, as predicted by our model.

**Fig. 5.**
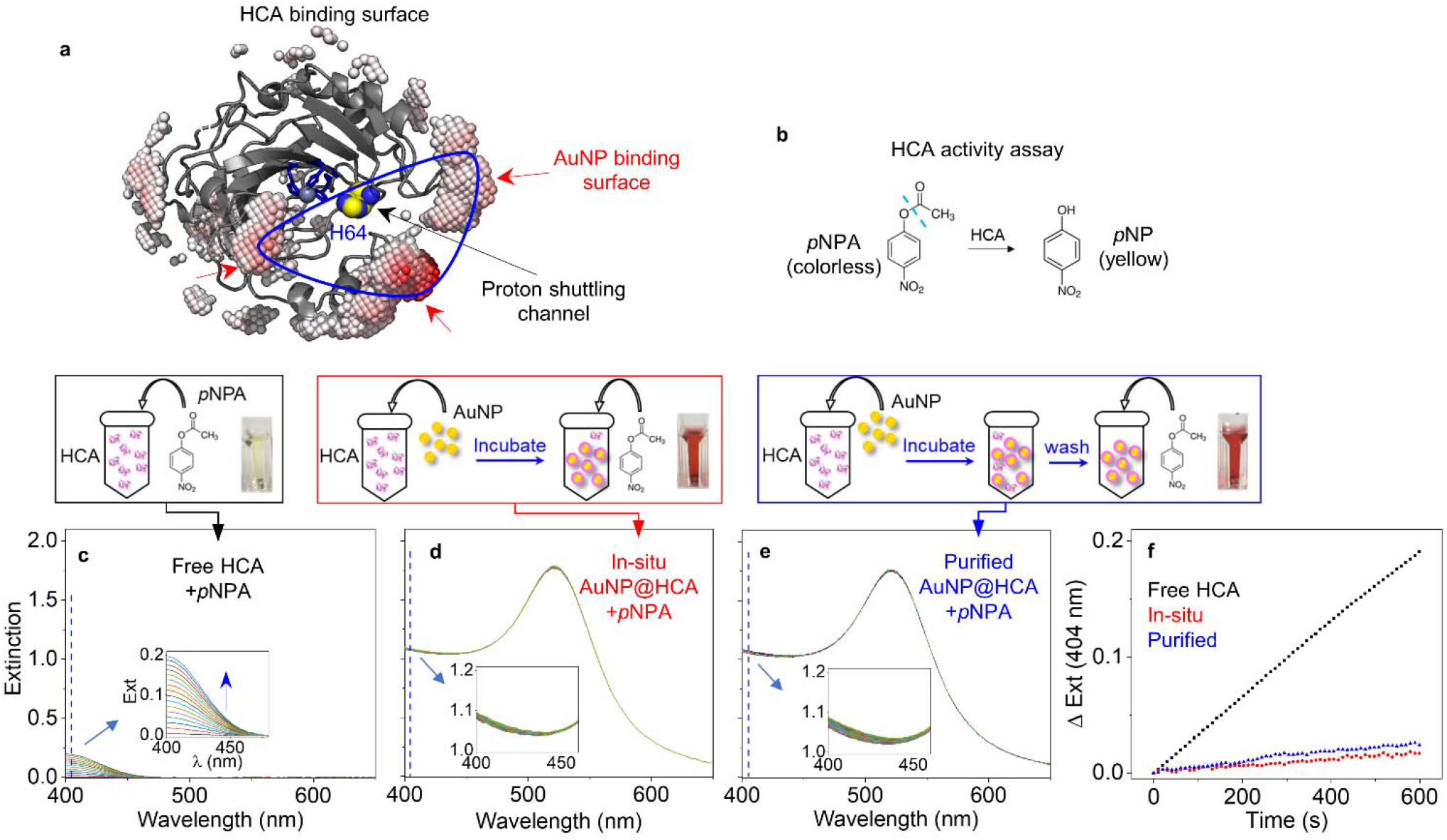
Prediction of bound orientation of human carbonic anhydrase (HCA) on AuNP. **a**, Cartoon structure of HCA showing the predicted AuNP-binding surface (highlighted blue region) with virtual atoms, with respect to its active center (highlighted in blue sticks), proton shuttling channel, Zn^2+^(gray sphere) and H64 (highlighted blue and yellow spheres). **b**, Mechanism of HCA assay of hydrolyzing colorless *p*NPA to yellow *p*NP, which absorbs at 404 nm. **c-e**, Time-resolved UV-vis spectra of HCA assay solutions with incubation time of 10 mins using free HCA (**c**), in-situ HCA bound to AuNPs (AuNP@HCA, **d**), and washed/purified AuNP@HCA (**e**). The enzyme/substrate ratio is fixed at 0.2 μM HCA/100 μM *p*NPA in all assays. The associated schemes above illustrate the sample preparation, and pictures show each solution after 30-min incubation. **f**, Comparisons of extinction change at 404 nm as a function of reaction time using free HCA (black), in-situ AuNP@HCA (red) and purified AuNP@HCA (blue). Close-up UV-Vis spectra are shown as insets in **c-e**.

Our approach is able to rationalize the enzyme behavior from the literature as well. No steric hindrance of the active site is predicted for two near-fully active AuNP-bound enzymes (acetylcholinesterase^37^ and citrate synthase^37^; all details, along with predicted orientations, are provided in the **Supplementary Information**). In addition, the predicted orientation of cytochrome c on citrate-reduced silver nanoparticles is consistent with observations by surface-enhanced Raman scattering^38^. In contrast, the activities of bound alcohol dehydrogenase^28^, horseradish peroxidase^39^ and glucose oxidase^40^ are significantly reduced on AuNPs. Our model successfully ascribes the behavior of alcohol dehydrogenase and horseradish peroxidase to partial blockage of the active site; however, the behavior of glucose oxidase is not predicted by our model. Lescovac *et al.* determined that glucose oxidase partially unfolds upon adsorption^40^, and our approach currently has no way to account for decreases in protein stability^41^. Overall, the effect of bound orientation on enzyme activities are successfully predicted by our model, but additional research is needed to understand energetics of protein-protein interactions and conformational changes that occur upon surface adsorption.

## Conclusions

In summary, we have determined an AuNP-binding affinity scale for each amino acid residue using a host-guest approach. Competition is an essential feature of this approach, because protein adsorption to citrate-AuNPs exhibits elements of both thermodynamic and kinetic control. We have demonstrated that this affinity scale can be used to map the residue-specific contribution to AuNP-binding for a protein, and therefore predict its bound orientation and function. This prediction is confirmed experimentally with two representative enzymes. This algorithm can also rationalize the known orientation of the GB3 protein on AuNPs, as well as five out of six other examples from the scientific literature. These results are significant given the biological relevance of protein’s orientation when bound to NPs. Until now, a straightforward algorithm to predict bound protein activity has not been available; such an algorithm should enable high-throughput predictions. Moving forward, this algorithm can potentially predict the composition of the protein corona formed in vivo by calculating the rank-order binding affinity of various blood proteins in serum. Such a prediction could have tremendous value in the development of nanomedicines with improved safety and functionality.

## Acknowledgements

We thank Joe Emerson, Julie Champion, and Dongmao Zhang for helpful suggestions in developing this project. This work was supported by the National Institute of Allergy and Infectious Diseases of the National Institutes of Health under grant number R01AI139479 and the National Science Foundation under grant number MCB 1818090. The content is solely the responsibility of the authors and does not necessarily represent the official views of the National Institutes of Health or National Science Foundation.

## Author contributions

J. X.X. and M.S.A. and created the initial GB3 variant library and synthesized all nanoparticles. M.S.A. assigned the NMR spectra, and performed initial nanoparticle binding experiments. J.X.X. performed competitive binding and titration experiments, designed and performed all experiments with protein prediction, and complied the figures. R.Y. assisted with NMR data acquisition and processing. N.C.F designed and directed the study and wrote python scripts. N.C.F and J.X.X analyzed the data and wrote the manuscript. All authors approved the final manuscript.

## Additional information

**Supplementary information** The online version contains supplementary material available at ….

## References

1. Lynch I., Dawson K. A. Protein-nanoparticle interactions. Nano Today 3, 40–47 (2008).

2. Milani S., Baldelli Bombelli F., Pitek A. S., Dawson K. A., Rädler J. Reversible versus irreversible binding of transferrin to polystyrene nanoparticles: Soft and hard corona. ACS Nano. 6, 2532–2541 (2012).

3. Liu J., Shi J., Nie W., Wang S., Liu G., Cai K. Recent progress in the development of multifunctional nanoplatform for precise tumor phototherapy. Adv. Healthc. Mater. 10, 2001207 (2021).

4. Cifuentes-Rius A., Desai A., Yuen D., Johnston A. P. R., Voelcker N. H. Inducing immune tolerance with dendritic cell-targeting nanomedicines. Nat. Nanotech. 16, 37–46 (2021).

5. Kelly P. M., Åberg C., Polo E., O’Connell A., Cookman J., Fallon J., et al. Mapping protein binding sites on the biomolecular corona of nanoparticles. Nat. Nanotech. 10, 472–479 (2015).

6. Mahon E., Salvati A., Baldelli Bombelli F., Lynch I., Dawson K. A. Designing the nanoparticle–biomolecule interface for “targeting and therapeutic delivery”. J. Controlled Release 161, 164–174 (2012).

7. Mahmoudi M., Lynch I., Ejtehadi M. R., Monopoli M. P., Bombelli F. B., Laurent S. Protein−nanoparticle interactions: Opportunities and challenges. Chem. Rev. 111, 5610–5637 (2011).

8. Monopoli M. P., Åberg C., Salvati A., Dawson K. A. Biomolecular coronas provide the biological identity of nanosized materials. Nat. Nanotech. 7, 779–786 (2012).

9. Zhang W., Besford Q. A., Christofferson A. J., Charchar P., Richardson J. J., Elbourne A., et al. Cobalt-directed assembly of antibodies onto metal–phenolic networks for enhanced particle targeting. Nano Lett. 20, 2660–2666 (2020).

10. Bednar R. M., Golbek T. W., Kean K. M., Brown W. J., Jana S., Baio J. E., et al. Immobilization of proteins with controlled load and orientation. ACS Appl. Mater. Interfaces 11, 36391–36398 (2019).

11. Lynch I., Cedervall T., Lundqvist M., Cabaleiro-Lago C., Linse S., Dawson K. A. The nanoparticle–protein complex as a biological entity; a complex fluids and surface science challenge for the 21st century. Adv. Colloid Interface Sci. 134-135, 167–174 (2007).

12. Yong K. W., Yuen D., Chen M. Z., Porter C. J. H., Johnston A. P. R. Pointing in the right direction: Controlling the orientation of proteins on nanoparticles improves targeting efficiency. Nano Lett. 19, 1827–1831 (2019).

13. Liu F., Wang L., Wang H., Yuan L., Li J., Brash J. L., et al. Modulating the activity of protein conjugated to gold nanoparticles by site-directed orientation and surface density of bound protein. ACS Appl. Mater. Interfaces 7, 3717–3724 (2015).

14. Jain A., Trindade G. F., Hicks J. M., Potts J. C., Rahman R., Hague R. J. M., et al. Modulating the biological function of protein by tailoring the adsorption orientation on nanoparticles. J. Colloid Interface Sci. 587, 150–161 (2021).

15. Zanzoni S., Pedroni M., D’Onofrio M., Speghini A., Assfalg M. Paramagnetic nanoparticles leave their mark on nuclear spins of transiently adsorbed proteins. J. Am. Chem. Soc. 138, 72–75 (2016).

16. Baldwin A. J., Kay L. E. NMR spectroscopy brings invisible protein states into focus. Nat. Chem. Biol. 5, 808–814 (2009).

17. Perera Y. R., Hill R. A., Fitzkee N. C. Protein interactions with nanoparticle surfaces: Highlighting solution NMR techniques. Isr. J. Chem. 59, 962–979 (2019).

18. Wang A., Perera Y. R., Davidson M. B., Fitzkee N. C. Electrostatic interactions and protein competition reveal a dynamic surface in gold nanoparticle–protein adsorption. J. Phys. Chem. C 120, 24231–24239 (2016).

19. Xu J. X., Alom M. S., Fitzkee N. C. Quantitative measurement of multiprotein nanoparticle interactions using NMR spectroscopy. Anal. Chem. 93, 11982–11990 (2021).

20. Xu J. X., Fitzkee N. C. Solution NMR of nanoparticles in serum: Protein competition influences binding thermodynamics and kinetics. Front. Physiol. 12, (2021).

21. Schöttler S., Landfester K., Mailänder V. Controlling the stealth effect of nanocarriers through understanding the protein corona. Angew. Chem. Int. Ed. 55, 8806–8815 (2016).

22. Vilanova O., Mittag J. J., Kelly P. M., Milani S., Dawson K. A., Rädler J. O., et al. Understanding the kinetics of protein–nanoparticle corona formation. ACS Nano. 10, 10842–10850 (2016).

23. Vangala K., Ameer F., Salomon G., Le V., Lewis E., Yu L., et al. Studying protein and gold nanoparticle interaction using organothiols as molecular probes. J. Phys. Chem. C 116, 3645–3652 (2012).

24. Wang A., Vangala K., Vo T., Zhang D., Fitzkee N. C. A three-step model for protein– gold nanoparticle adsorption. J. Phys. Chem. C 118, 8134–8142 (2014).

25. Inkpen M. S., Liu Z. F., Li H., Campos L. M., Neaton J. B., Venkataraman L. Non-chemisorbed gold–sulfur binding prevails in self-assembled monolayers. Nat. Chem. 11, 351–358 (2019).

26. Zamyatnin A. A. Protein volume in solution. Prog. Biophys. Mol. Biol. 24, 107–123 (1972).

27. Ly N. H., Oh C. H., Joo S.-W. A submicromolar Cr(III) sensor with a complex of methionine using gold nanoparticles. Sensors Actuators B: Chem. 219, 276–282 (2015).

28. Petkova G. A., Záruba К., Žvátora P., Král V. Gold and silver nanoparticles for biomolecule immobilization and enzymatic catalysis. Nanoscale Res. Lett. 7, 287 (2012).

29. Lacerda S. H. D. P., Park J. J., Meuse C., Pristinski D., Becker M. L., Karim A., et al. Interaction of gold nanoparticles with common human blood proteins. ACS Nano. 4, 365–379 (2010).

30. Monopoli M. P., Walczyk D., Campbell A., Elia G., Lynch I., Baldelli Bombelli F., et al. Physical−chemical aspects of protein corona: Relevance to in vitro and in vivo biological impacts of nanoparticles. J. Am. Chem. Soc. 133, 2525–2534 (2011).

31. Hubbard S., Thornton J. Naccess: Program for calculating accessibilities. Department of Biochemistry and Molecular Biology, University College of London, (1992).

32. Chan K. P., Chao S.-H., Kah J. C. Y. Exploiting protein corona around gold nanoparticles conjugated to p53 activating peptides to increase the level of stable p53 proteins in cells. Bioconj. Chem. 30, 920–930 (2019).

33. Biscaglia F., Caligiuri I., Rizzolio F., Ripani G., Palleschi A., Meneghetti M., et al. Protection against proteolysis of a targeting peptide on gold nanostructures. Nanoscale 13, 10544–10554 (2021).

34. Krishnamurthy V. M., Kaufman G. K., Urbach A. R., Gitlin I., Gudiksen K. L., Weibel D. B., et al. Carbonic anhydrase as a model for biophysical and physical-organic studies of proteins and protein−ligand binding. Chem. Rev. 108, 946–1051 (2008).

35. Fisher S. Z., Maupin C. M., Budayova-Spano M., Govindasamy L., Tu C., Agbandje-McKenna M., et al. Atomic crystal and molecular dynamics simulation structures of human carbonic anhydrase II:LJ Insights into the proton transfer mechanism. Biochemistry 46, 2930–2937 (2007).

36. Saada M.-C., Montero J.-L., Vullo D., Scozzafava A., Winum J.-Y., Supuran C. T. Carbonic anhydrase activators: Gold nanoparticles coated with derivatized histamine, histidine, and carnosine show enhanced activatory effects on several mammalian isoforms. J. Med. Chem. 54, 1170–1177 (2011).

37. Baruah P., Yesylevskyy S. O., Aguan K., Mitra S. Modulation of enzyme activity at nano-bio interface: A case study with acetylcholinesterase and citrate synthase adsorbed on colloidal metal nanoparticles. J. Mol. Liq. 325, 115201 (2021).

38. Macdonald I. D. G., Smith W. E. Orientation of cytochrome c adsorbed on a citrate-reduced silver colloid surface. Langmuir 12, 706–713 (1996).

39. Cans A. S., Dean S. L., Reyes F. E., Keating C. D. Synthesis and characterization of enzyme-Au bioconjugates: HRP and fluorescein-labeled HRP. NanoBiotechnology 3, 12–22 (2007).

40. Leskovac V., Trivić S., Wohlfahrt G., Kandrač J., Peričin D. Glucose oxidase from *Aspergillus niger*: The mechanism of action with molecular oxygen, quinones, and one-electron acceptors. Int. J. Biochem. Cell Biol. 37, 731–750 (2005).

41. Woods K. E., Perera Y. R., Davidson M. B., Wilks C. A., Yadav D. K., Fitzkee N. C. Understanding protein structure deformation on the surface of gold nanoparticles of varying size. J. Phys. Chem. C 120, 27944–27953 (2016).

