## Supplementary material for "Predicting Protein Function and Orientation on a Gold Nanoparticle Surface Using a Residue-Based Affinity Scale": Supplrementary Information

### Additional Supplementary Methods

#### Chemicals and Materials

Gold (III) chloride trihydrate (product #: 520918) and sodium citrate dihydrate (product #: 567446) were purchased from Millipore Sigma. The NMR reference compound deuterated 3-1-propanesulfonic acid-d6 sodium salt (DSS-d6, DLM-8206), Tryptophan ($^{15}$N labeled, NLM-800-PK), and 99.9% $D_2O$ (DLM-4) were purchased from Cambridge Isotope Labs (Tewksbury, MA). All chemicals were used as purchased without additional purification. Proteinase K (PK) was obtained from Amresco (#0706, biotechnology grade). Bovine serum albumin (BSA) was purchased from Calbiochem (# 12657, electrophoresis 100%). Human carbonic anhydrase II (HCA) was provided by Dr. Joseph P. Emerson at Mississippi State University. Before use, HCA was converted to apoCA by dialysis with ACES buffer containing dipicolinic acid at pH 7 described previously.[1] *p*-Nitrophenyl acetate (*p*NPA) was purchased from Sigma-Aldrich (#N8130).

#### Synthesis and characterizations of 15-nm AuNPs.

The method developed by Frens and Turkevich was employed to synthesize AuNPs.[2, 3] All glassware was cleaned with aqua regia and thoroughly rinsed with ultrapure water prior to each use. Briefly, 2 mL of 10 mg $mL^{-1}$ $HAuCl_4$ was added into 198 mL of 18.2 MΩ ultrapure Milli-Q water in a round bottom flask equipped with a condenser. The reaction was constantly stirred at 750 rpm and heated to boil at power level 70 in a heating mantle (Glas-Col). Once the boiling started, the power level of the mantle was set to zero. When the boiling stopped with no bubbling, 6.5 ml of 1% (w/v) sodium citrate was added, and the power level of the heating mantle was set to 48. The solution was heated at this new power level for 25 minutes. Subsequently, the flask was removed from the heating mantle and allowed to cool down with constant stirring. When the reaction reached room temperature, 800 µL of 0.5 M sodium citrate was added to stabilize the newly synthesized AuNPs. The synthesized AuNPs were characterized by DLS and UV-vis spectrometry (**Supplementary Fig. 1**) to ensure their sizes are 15 nm with polydispersity within 10%. Prior to preparation of NMR samples, the AuNP solution was centrifuged in 50 mL Falcon tubes at 8,000 *g* for 45 minutes at 15 °C. Afterward, the supernatant was removed carefully to concentrate the AuNPs from ~2 nM to 200-300 nM. The exact AuNP concentration was measured by diluting 1 µL of the concentrated AuNP solution into a 1000 µL solution, and determined by its UV-vis peak absorption at 520 nm using a molar absorptivity of $3.94 \times 10^8$ $M^{-1}cm^{-1}$.[2]

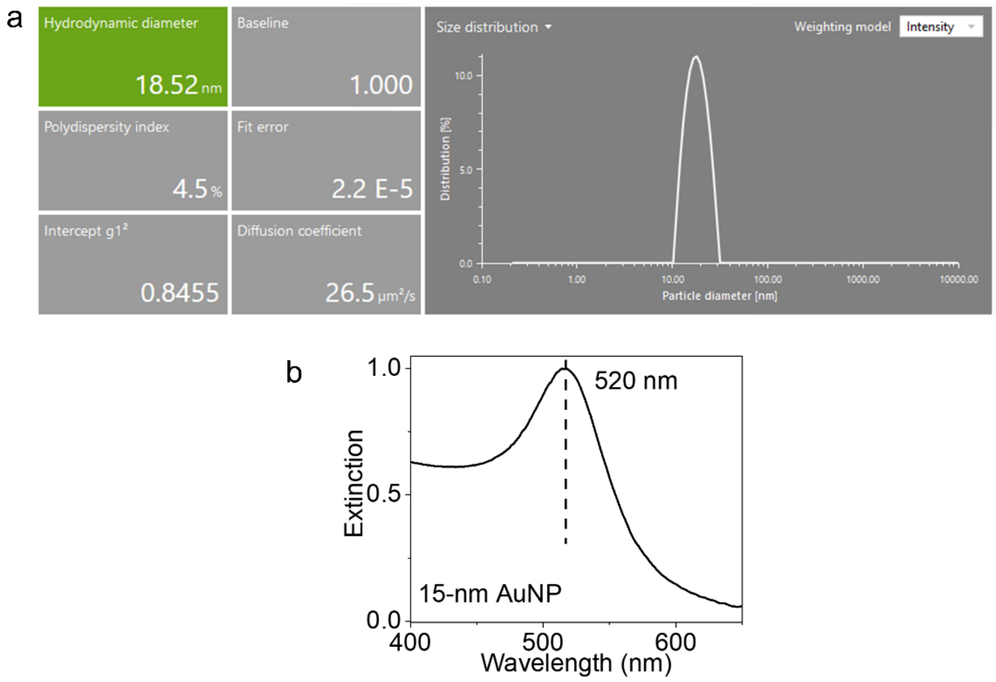

**Supplementary Fig. 1 | Characterization of as-synthesized 15-nm citrate-capped AuNPs. a,** Particle size distribution determined by Anton Paar DLS system. **b,** UV-vis extinction spectrum of as-synthesized AuNPs.

### Site-directed mutagenesis for 20 GB3 variants

Site-directed mutagenesis was performed on the pGS-21 vector of the wild-type (wt) GB3 sequence (**Supplementary Fig. 2**) to introduce K13X point mutation.

```
                                M  Q  Y  K  L  V  I  N  G  K  T  L  K    13
1    ctttaagaaggagatatacatatgcagtacaaattagttatcaatggtaaaacattgaaa  60

14   G  E  T  T  T  K  A  V  D  A  E  T  A  E  K  A  F  K  Q  Y    33
61   ggcgaaacaactactaaagctgttgatgctgaaactgcagaaaaagctttcaaacaatac  120

34   A  N  D  N  G  V  D  G  V  W  T  Y  D  D  A  T  K  T  F  T    53
121  gctaacgacaacggtgttgacggtgtttggacttacgacgatgcgactaagacctttaca  180

54   V  T  E  *
181  gttactgaataggatccggctgctaacaaagcc  213
```

**Supplementary Fig. 2 |** DNA and translated protein sequence of wt GB3. In this figure, the highlighted, bold M represents the start of translation, and the asterisk represents the stop codon. One-letter amino acid codes are shown on the top line in capital letters, and the DNA sequence is shown in the lower line in lower case. Flanking, untranslated nucleotides from the pGS-21 vector are shown in addition to the translated sequence.

Primers were designed and optimized using GeneRunner Software version 6.5.52 (http://www.generunner.net/) and the New England Biolabs (NEB) $T_m$ Calculator (https://tmcalculator.neb.com/#!/main). Parameters including melting temperature, percent of GC content, annealing temperature ($T_m$), probability of primer dimerization, and the number of nucleotides were optimized. A total of 19 sets of primers used in this study are listed in **Supplementary Table 1**. The bases in red represent the codons that were changed. However, a single nucleotide in the codon of K10 (AAA to AAG) was also changed for K13F, K13N, K13Y, and K13M mutants. This change does not alter K10, but it increases the GC content and was found to increase the polymerase chain reaction (PCR) success rate.

Site-directed Mutagenesis was performed with PCR using the Phusion High-Fidelity DNA Polymerase kit by NEB (NEB # E0553S). Stock solutions of 10 µM forward (Fwd) and reverse

(Rev) primers were made using 18.2 MΩ ultrapure Milli-Q water. Wild type pGS-21 plasmid was extracted from a 10 ml culture media of *E. Coli* XL1 Blue (Invitrogen) cells using a miniprep plasmid extraction protocol,[3] and a 20 ng/µL stock solution was obtained as the DNA template for PCR. In a typical PCR setup, final concentrations of 0.5 uM Fwd primer, 0.5 uM Rev primer, 200 uM dNTPs, 20 ng template DNA, 1.0 unit of Phusion DNA polymerase were used. Detailed mixing protocols are presented in **Supplementary Table 2**. Subsequently, the reaction was carried out in a PCR thermocycler (GeneAmp PCR system 9700). The upper thermal blocks of a PCR machine were preheated to 98°C before transferring PCR tubes from ice to prevent condensation on the cap during heating. Thermocycling conditions for the PCR used in this work are given in **Supplementary Table 3**. The denaturation, annealing, and elongation steps were repeated for 28 cycles.

The PCR product was analyzed on DNA gel. Briefly, 10 uL of PCR product was stained un a gel containing a final concentration of 0.5 ug/mL Ethidium Bromide (EtBr). A 0.5% agarose (Sigma Aldrich) gel was cast, and electrophoresis was performed at a constant 75 V for 90 minutes. The gel was cast using 1 X TAE buffer with 0.5µg/mL EtBr. Once electrophoresis was finished, the gel was carefully transferred to a UV-photographic system for visualizing the bands corresponding to plasmid lengths compared with the standard DNA ladder.

The template DNA was digested by DpnI reaction using the DpnI kit (NEB #R0176L). 1uL of the DpnI restriction enzyme was added to the remaining PCR product (~40 µL after gel analysis), 5 µL NEBuffer, and MQ water to make a 50 µL reaction. Subsequently, the reaction mixture was incubated at 37 $^{o}$C in the PCR thermocycler for an hour. A PCR cleanup kit (Wizard SV, Promega) was then used to purify the synthesized DNA after the DpnI reaction.

The purified DNA was transformed to competent *E. Coli* XL1 Blue cells by heat shock DNA transformation.[4] These cells were plated on ampicillin-containing LB media and grown overnight at 37℃. Single colonies from the plates were transferred to ampicillin containing lysogeny broth (LB) media for overnight growth, and plasmids were extracted using the miniprep plasmid extraction protocol (Wizard SV, Promega).[3] To get the final confirmation of a successful mutation, 10 µL purified plasmid was sent for Sanger sequencing (Eurofins Genomics) using the T7 promoter.

**Supplementary Table 1 | PCR Primers used in this study**

| Primer Name | Primer Sequence (5' to 3') |
|---|---|
| K13R -Fwd | GGTAAAACATTGCGTGGCGAAACAACTACTAAAGCTG |
| K13R -Rev | GCTTTAGTAGTTGTTTCGCCACGCAATGTTTTACCATTG |
| K13D -Fwd | GGTAAAACATTGGACGGCGAAACAACTACTAAAGCTG |
| K13D -Rev | GCTTTAGTAGTTGTTTCGCCGTCCAATGTTTTACCATTG |
| K13E -Fwd | GGTAAAACATTGGAGGGCGAAACAACTACTAAAGCTG |
| K13E -Rev | GCTTTAGTAGTTGTTTCGCCCTCCAATGTTTTACCATTG |
| K13T -Fwd | GGTAAAACATTGACGGGCGAAACAACTACTAAAGCTG |
| K13T -Rev | GCTTTAGTAGTTGTTTCGCCCGTCAATGTTTTACCATTG |
| K13V -Fwd | GGTAAAACATTGGTGGGCGAAACAACTACTAAAGCTG |
| K13V -Rev | GCTTTAGTAGTTGTTTCGCCCACCAATGTTTTACCATTG |
| K13C -Fwd | GGTAAAACATTGTGCGGCGAAACAACTACTAAAGCTG |
| K13C -Rev | GCTTTAGTAGTTGTTTCGCCGCACAATGTTTTACCATTG |
| K13G -Fwd | GGTAAAACATTGGGTGGCGAAACAACTACTAAAGCTG |
| K13G -Rev | GCTTTAGTAGTTGTTTCGCCACCCAATGTTTTACCATTG |
| K13I -Fwd | GGTAAAACATTGATCGGCGAAACAACTACTAAAGCTG |
| K13I -Rev | GCTTTAGTAGTTGTTTCGCCGATCAATGTTTTACCATTG |
| K13F -Fwd | GGTAAGACATTGTTCGGCGAAACAACTACTAAAGCTG |

| Primer Name | Primer Sequence (5' to 3') |
|---|---|
| K13F -Rev | GCTTTAGTAGTTGTTTCGCCGAACAATGTCTTACCATTG |
| K13Y -Fwd | GGTAAGACATTGTACGGCGAAACAACTACTAAAGCTG |
| K13Y -Rev | GCTTTAGTAGTTGTTTCGCCGTACAATGTCTTACCATTG |
| K13N -Fwd | GGTAAGACATTGAACGGCGAAACAACTACTAAAGCTG |
| K13N - Rev | GCTTTAGTAGTTGTTTCGCCGTTCAATGTCTTACCATTG |
| K13M -Fwd | GGTAAGACATTGATGGGCGAAACAACTACTAAAGCTG |
| K13M -Rev | GCTTTAGTAGTTGTTTCGCCCATCAATGTCTTACCATTG |
| K13A -Fwd | GGTAAAACATTGGCAGGCGAAACAACTACTAAAGCTG |
| K13A -Rev | GCTTTAGTAGTTGTTTCGCCTGCCAATGTTTTACCATTG |
| K13L -Fwd | GGTAAAACATTGCTGGGCGAAACAACTACTAAAGCTG |
| K13L -Rev | GCTTTAGTAGTTGTTTCGCCCAGCAATGTTTTACCATTG |
| K13S -Fwd | CATTGAGCGGCGAAACAACTAC |
| K13S -Rev | CAAAGCGGCGCTTTACAAAATG |
| K13H -Fwd | CATTGCACGGCGAAACAACTAC |
| K13H -Rev | CAAAGCGGCGTGTTACAAAATG |
| K13W-Fwd | CATTGTGGGGCGAAACAACTAC |
| K13W -Rev | CAAAGCGGCCCATTACAAAATG |
| K13P-Fwd | CATTGCCGGGCGAAACAACTAC |
| K13P -Rev | CAACAAAGCGGCCGGTTACAAAATG |
| K13Q-Fwd | CATTGCAGGGCGAAACAACTAC |
| K13Q -Rev | CAACAAAGCGGCCTGTTACAAAATG |

**Supplementary Table 2 | Typical PCR reaction preparation**

| Component | Stock Solution | Volume Added (μl) |
| --- | --- | --- |
| dNTPs | 10 mM | 1.0 |
| Forward Primer | 10 μM | 2.5 |
| Reverse Primer | 10 μM | 2.5 |
| Template DNA | 20 ng/μL | 1.0 |
| HF Buffer | 5 X | |
| Ultrapure Water | - | 32.5 |
| Phusion DNA Polymerase | - | 0.5 |
| **Total Volume** | **-** | **50** |

**Supplementary Table 3 | Thermocycling conditions**

| Steps | Temperature | Time |
| --- | --- | --- |
| Initial Denaturation | 98℃ | 30 seconds |
| Denaturation | 98℃ | 30 seconds |
| Primer Annealing ($T_m$) | 54℃ | 20 seconds |
| Elongation | 72℃ | 4 minutes |
| Final Elongation | 72℃ | 10 minutes |
| Hold | 4℃ | - |

**Expression and Purification of GB3 Variants**

The mutant DNA plasmids extracted from XL1 Blue cells were transformed into BL21(Star) DE3 for protein expression. Each variant was expressed and purified by employing the Gräslund Method.[5]

Briefly, the mutant Bl21(Star) DE3 cells (Invitrogen) and grown for 6 hours in 5 mL ampicillin containing LB medium were inoculated into 25mL ampicillin-containing $^{15}N$ M9 media (starter culture) and grown overnight at 37°C. The starter culture was then transferred into a 1L $^{15}N$ M9 media to reach an initial $OD_{600}$ of 0.05. The culture flasks were incubated in a shaker at 37°C and at 200 rpm and induced with 0.5ml of 1mM isopropyl β,D-1-thiogalactopyranoside (IPTG) when the $OD_{600}$ reached 0.5-0.7. After induction, cells were allowed to express protein for approximately 5 hours at 37°C temperature, and $OD_{600}$ was monitored every hour. Cells were harvested by centrifugation at 7,000 *g* for 20 min and cell pellets were collected for the next step (or stored at -80°C if not used immediately). The harvested cells were resuspended in 20 mL cold lysis buffer (50 mM NaCl, 20 mM $NaH_2PO_4$ pH 4.5, 5 mM EDTA, 0.5 mg/mL lysozyme) on ice. The resuspended cells were sonicated on ice at 45% power for 6 minutes of total processing time (30 seconds pulse, 30 seconds rest). After lysis, the lysed cell solution was incubated in an 85°C hot water bath for 15 minutes to allow other cellular proteins to unfold and precipitate, with swirling every 3-4 minutes. Afterward, the solution was immediately transferred to an ice water bath, and streptomycin sulfate was added (0.5 mg/mL final concentration) to precipitate DNA. After 15 minutes on ice, denatured proteins and DNA were removed by centrifugation at 18,000 *g* for 45 minutes. The supernatant was collected and loaded on an AKTA purification system using a 5 mL HiTrap Q FF anion exchange column for purification. Under these conditions (pH 4.5), the column binds additional DNA impurities, while GB3 is flows through in the wash. Fractions with high UV-vis absorption at 280 nm were collected and pooled. The collected fractions were concentrated using 3.0 kDa size EMD Millipore Amicon Ultra Centrifugal Filters (Pall Corporation). For K13C GB3, a final concentration of 5 mM DTT was added to all buffers.

The concentrated protein was further loaded to a HiLoad 26/600 Superdex 75pg gel filtration column equilibrated with gel filtration buffer (50 mM NaCl, 20 mM $NaH_2PO_4$ pH 4.5). The fractions of K13X GB3 were found to elute around 210 mL with a stronger absorbance at 280 nm than 260 nm. The purified K13X GB3 protein was dialyzed in 10 mM HEPES buffer at pH 6.5 and 5 mM NaCl for 5 hours (plus 1 mM TCEP for K13C). All GB3 variants were analyzed by gel electrophoresis to ensure purity.

**Characterization of GB3 variants by 2D TOCSY-HSQC NMR.**

The traditional HSQC, NOESY-HSQC,[4] and TOCSY-HSQC [5] experiments were performed to assign and backbone amide peaks. A 500 μM $^{15}N$ K13X GB3 protein sample was prepared in HEPES at pH 6.5 and 6% $D_2O$ for the measurement (with 1 mM TCEP for K13C GB3). $^{1}H$-$^{15}N$ HSQC reveals the correlation of the backbone $^{15}N$ and its amide proton of each residue. TOCSY and NOESY spectra were sufficient to assign variants of WT GB3. As the example illustrated in **Supplementary Fig. 3a**, the amide proton NOE from the preceding T11 amide appears in the $^{N}H$ strip of L12, and the Hα NOE of L12 shows up in the strip for H13. A comparison of $^{1}H$-$^{15}N$ HSQC spectra of wt GB3, K13Q, and K13S is presented in **Supplementary Fig. 3b,** which shows K13 at 123.8 ppm, Q13 at 122.8 ppm, and S13 at 117.0 ppm. The vast majority of GB3's 56 backbone amide peaks exhibit only a marginal chemical shift perturbation when position 13 is changed; this demonstrates that mutagenesis does not significantly change the folded structure of GB3. The $^{15}N$ and $^{1}H$ chemical shifts of each variant are presented in **Supplementary Tables S6-S12**.

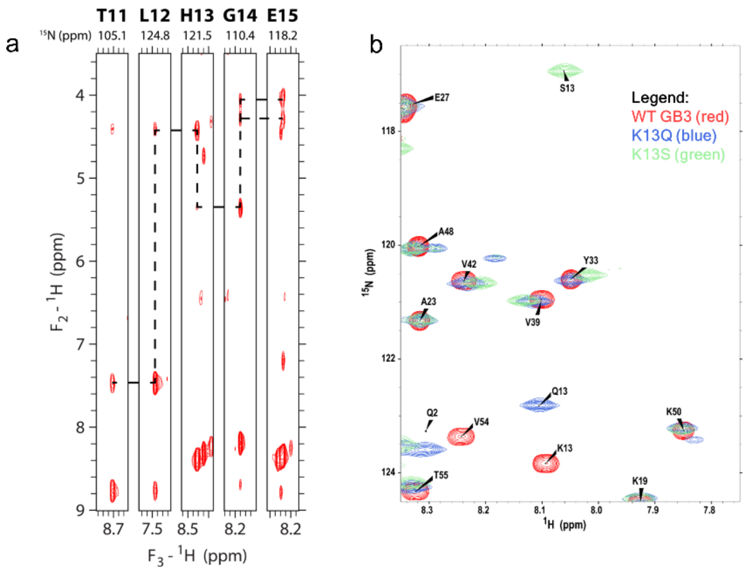

**Supplementary Fig. 3 | a,** NOESY strip plot identifies assignment of residue 13 of K13H GB3 using the alpha proton connectivity. **b,** HSQC spectra comparing wt GB3 and two K13X variants. Differences in line widths result from differing acquisition times and do not reflect changes in protein motions.

### Preparation of competitive binding samples of two GB3 variants

Stock solutions of ~150 µM GB3 variants, 100 mM HEPES buffer at 6.5, 500 mM NaCl solution, and ~300 nM AuNP solution were prepared. The extinction coefficient for each variant was determined using the approach of Pace, *et al*.[5] To set up competitive binding experiments for each variant, a protein/AuNP mixture (P/NP) NMR sample of a total volume of 400 µL was made with 20 µM $^{13}$C K13G GB3, 20 µM $^{15}$N K13X GB3, 20 mM HEPES buffer at pH 6.5, 20 mM NaCl, 18.2 MΩ ultrapure Milli-Q water and 50 nM AuNPs. A protein (P) sample as control was made identically except without the addition of AuNPs. All binding experiments were conducted under pH 6.5 unless indicated otherwise. 100 mM NaOAc/AcOH buffer at pH 5.0 and 100 mM HEPES at pH 8.0 are used instead for pH-dependent binding experiments. The protein/AuNP sample was incubated for an hour before NMR experiment to ensure the binding had reached equilibrium. 4 mM 4,4-dimethyl-4-silapentane-1-sulfonic acid (DSS) was used as an external standard for peak intensity calibration.

**Affinity scale for residue X quantified by 1D filtered NMR experiments**

Proton 1D NMR spectra were recorded on a 600 MHz Bruker Avance III cryoprobe-equipped NMR spectrometer with a jump-return program for water suppression.[6] An acquisition time of 80 ms was used, and the inter-pulse delay of 110 μs was selected for proton resonances at 8.5 ppm. The total experiment time was 9 mins. Three 1D spectra were collected for each sample: A non-filtered experiment for calibrating peak intensities of the DSS reference between the Protein/NP and Protein samples; A $^{13}$C filtered experiment to obtain the signals from $^{13}$C K13G GB3 only; A $^{15}$N filtered experiment to measure the peak intensities solely from $^{15}$N K13X GB3 (**Figure 1, main text**). The binding of GB3 onto AuNP has proven to be in slow exchange regime, and therefore the NMR signal of bound proteins will be invisible due to peak broadening and the remaining signal is attributed to the unbound protein. By comparing the protein NMR signal without (P sample) and with (P/NP sample) the presence of AuNP, the bound amount of $^{13}$C K13G GB3 and $^{15}$N K13X GB3 can be determined, respectively. The data processing was performed with TOPSPIN. All 1D NMR spectra are phase adjusted identically and baseline subtracted. The DSS peak intensity was used as a reference to normalize all NMR spectra before comparison. The "scale spectrum" in the TOPSPIN was then used to determine the percentage of integrated proton NMR signal has dropped after the addition of AuNPs, which corresponds to the bound percentages of each variant (**Figure 1, main text**). Finally, the alpha value ($\alpha$) of K13X GB3 over K13G GB3 was quantified using Equation 1. An $\alpha$ value larger than 1 indicates K13X GB3 binds stronger than K13G GB3 when both are present competing for limited AuNP surface, and vice versa.

$$\alpha = \frac{[\,^{15}N\ K13X]_{bound}}{[^{13}C\ K13G]_{bound}} \quad \text{(Eq 1)}$$

where $\alpha_X$ is the affinity for residue X, $[\,^{15}N\ K13X]_{bound}$ and $[^{13}C\ K13G]_{bound}$ represent the bound concentrations of $^{15}$N-K13X GB3 and $^{13}$C-K13G GB3, respectively.

All $\alpha$ values are presented in **Table 1, main text**. The errors for all variants are identically represented by an average of 95% confidence intervals of 0.04 obtained from alpha values of 5 random variants with three observations. Each of these five variants were measured three times using independent replicate experiments. All other variants' alpha values were measured twice (**Supplementary Table 4**). The uncertainty for every GB3 variant should be identical, because all experiments are prepared and measured in the same way except for the variant of interest. The table below shows that the average 95% confidence interval of 0.04 is a reasonable upper limit for ascertaining alpha value differences. Moreover, all of the samples where two only two trials are performed are consistent with this confidence interval. The 95% confidence interval is calculated based on $\sigma = t * \frac{s}{\sqrt{n}}$ using the Excel built-in function of CONFIDENCE.T.

**Supplementary Table 4 |** Experimental replicates of alpha values and estimation of uncertainty

| Variant | Alpha Trial 1 | Alpha Trial 2 | Alpha Trial 3 | Sample Standard Deviation (*s*) | Average Alpha | 95% CI |
|---|---|---|---|---|---|---|
| K13G | 0.960 | 1.000 | | 0.029 | 0.980 | |
| K13A | 0.652 | 0.668 | 0.661 | 0.008 | 0.660 | 0.019 |
| K13L | 0.432 | 0.368 | | 0.045 | 0.400 | |
| K13I | 0.389 | 0.415 | | 0.018 | 0.402 | |
| K13V | 0.445 | 0.462 | 0.459 | 0.009 | 0.455 | 0.022 |
| K13M | 0.723 | 0.764 | | 0.029 | 0.743 | |
| K13P | 0.615 | 0.635 | | 0.015 | 0.625 | |
| K13F | 0.448 | 0.474 | | 0.019 | 0.461 | |
| K13Y | 0.463 | 0.470 | 0.489 | 0.014 | 0.474 | 0.034 |
| K13W | 0.401 | 0.423 | | 0.016 | 0.412 | |
| K13S | 0.570 | 0.570 | | 0.000 | 0.570 | |
| K13C | 5.847 | 5.536 | | 0.220 | 5.692 | |
| K13T | 0.506 | 0.497 | | 0.007 | 0.502 | |
| K13E | 0.374 | 0.356 | 0.324 | 0.025 | 0.352 | 0.063 |
| K13D | 0.488 | 0.445 | | 0.030 | 0.466 | |
| K13N | 0.529 | 0.553 | | 0.016 | 0.541 | |
| K13Q | 0.400 | 0.403 | | 0.002 | 0.401 | |
| K13H | 0.816 | 0.766 | | 0.035 | 0.791 | |
| K13K | 0.805 | 0.795 | 0.838 | 0.023 | 0.812 | 0.056 |
| K13R | 0.849 | 0.825 | | 0.017 | 0.837 | |
| | | | | **Average Confidence Interval** | | **0.04** |

**UV-vis titration for thermodynamic characterization of GB3 adsorption on AuNP**

UV-vis titrations were performed on 5 selective GB3 variants with an Olis-refurbished Agilent 8453 spectrophotometer. A series of 15 titration samples were prepared by adding protein with increasing concentrations (0-1600 nM) into 2.0 nM AuNP solutions in 20 mM HEPES (pH 6.5). UV-vis measurement was conducted after 1 h sample incubation. Subsequently, the maximum AuNP plasmonic peak wavelength for each sample was interpolated to a precision of 0.05 nm, using polynomial fitting with an order of 8.

**Binding kinetics of GB3 onto AuNP monitored by SOFAST-HMQC.**

The kinetics of individual or competitive binding of GB3 variants were monitored by SOFAST-HMQC 2D NMR (Bruker parameter set: sfhmqcf3gpphiasi).[7, 8] The parameters used include 32 points in the indirect dimension, acquisition time of 15 ms, 16 scans, and a recycle delay of 100 ms. The samples were prepared as follows: 20 µM K13X GB3 is prepared in 20 mM HEPES buffer and 20 mM NaCl as the Protein sample for "0 min point". The Protein/NP sample for kinetics measurement was prepared identically as the Protein-only sample, but with addition of 50 nM AuNP. Immediately after mixing the protein and nanoparticle solutions, the Protein/NP sample was vortexed and transferred to an NMR tube with the $^{15}$N-Trp reference insert,[9] and the tube was loaded into the NMR for shimming and measurement. The Protein-only sample was used for 3D shimming, while 1D-shimming was used for the Protein/NP due to its time sensitivity. Including mixing and shimming, the dead time of the experiment is approximately 5 mins before spectral acquisition of the Protein/NP sample. The acquisition of one SOFAST-HMQC spectrum is around 5 mins. A series of SOFAST-HMQC spectra were taken every 5 mins for a total time of 365 mins.

**Surface prediction for AuNP binding using alpha values**

First, the PDB file of a protein of interest (e.g., pepsin, 3PEP) is downloaded and opened with PyMOL (Schrodinger). Water molecules, extra chains, and hetero atoms are removed with PyMOL and the "clean" protein structure is saved as a new PDB file (`3pep_processed.pdb`). The calculation of binding affinity for each residue is completed by an in-house python script "asa_alpha_all-bfac.py" below. This script, and all others, are maintained at the Fitzkee Lab GitHub repository (https://github.com/FitzkeeLab/citrate-aunp-predict). Briefly, the binding affinity of each residue is calculated as a multiplication product of α values (ALPHA) and relative side chain accessible surface area (RASA). RASA was obtained by running NACCESS,[10] and only RASA larger than 25% was used. A new PDB file (`3pep_alpha.pdb`) is created with the B-factor column changed into product. The command used line is:

```
python3 ./asa-alpha_all_bfac.py 3pep_processed.pdb 3pep_alpha.pdb
```

To visualize the binding affinity with the use of "virtual atoms", we ran another in-house python script "binding_surface.py" on the modified PDB file (`3pep_alpha.pdb`) generated from the last step. In this processing script, the protein is placed on a $1 \times 1 \times 1$ angstrom grid, and only the grid points that are more than two angstroms, but less than three angstroms from protein atoms are selected. This selects only the grid points on the surface and some cavities. A virtual "atom" is placed at each grid point. The B-factor of for each virtual atom is calculated as the average product ($\alpha \times RASA$) of all protein atoms within 10 angstroms of the grid point. Only the grid points with an average product greater than 30 are written in a new PDB file. The output will list the maximum b-factor and the x, y, z coordinate of the largest b-factor. The command line to run this script is:

```
python3 ./binding_surface.py 3pep_alpha.pdb 3pep_mesh.pdb
```

Finally, the structure can be visualized using `3pep_alpha.pdb` and `3pep_mesh.pdb` in PyMOL. The following commands to depict the average values from white to red, to indicating low vs. high binding affinity, as presented in the text.

```
show spheres, 3pep_mesh
spectrum b, white_red, 3pep_mesh, minimum=30, maximum=50,
selection=3pep_mesh
```

```
set sphere_scale, 0.5, 3pep_mesh
```

Additional instructions are included on the GitHub site.

**Proteinase K proteolytic activity assay**

Proteinase K (PK) was dissolved in 10 mM $KH_2PO_4$ at pH 7.5 and applied to a desalting column (Thermo Scientific, # 89882) before use. Its concentration (mg/mL) was determined by UV absorption at 280 nm ($\varepsilon_{280} = 36{,}580\ M^{-1}cm^{-1}$) and molecular weight (M.W.= 28,907 g/mol). The activity of PK was evaluated by its proteolytic reaction with BSA as the substrate. Different PK/BSA ratios were explored for incomplete BSA digestion, and 0.01 mg/mL PK and 2 mg/mL BSA was finally used for the assay condition.

In order to control for the concentration of AuNP-bound PK in the proteolytic reaction, the binding capacity of PK on 15-nm AuNP was quantified by 1D proton NMR.[6] 50 nM of AuNP was mixed with 20 µM PK in 10 mM $KH_2PO_4$ at pH 7.5, the proton intensities of which are compared with those from 20 µM PK without addition of AuNPs. A scale factor (0.795 ± 0.014) was obtained using TOPSPIN, which corresponds to the unbound fraction of PK. The binding capacity of PK ($C_{PK}$) was determined as 82 ± 6 PK per AuNP (**Supplementary Fig. 4**). The AuNP-bound PK was prepared in two ways, in situ (in-situ AuNP@PK) and by washing (purified AuNP@PK). For the in situ method, based on its $C_{PK}$, 6.8 nM AuNP was mixed with 0.01 mg/mL PK for at least three hours to ensure that ~90% PK proteins are tightly bound before adding BSA. We make sure all AuNP surfaces are saturated (leaving ~10% excess PK unbound) to avoid BSA binding to AuNP, which would effectively protect BSA from proteolysis, as discussed in the main text. For the washing method, an excess amount of 20 µM PK was mixed with 50 nM AuNPs for three hours to reach binding equilibrium (solution volume of 400 uL), and the PK-coated AuNPs (AuNP@PK) was spun down by centrifugation at 9,000 $g$ for 15 mins. The supernatant was carefully removed, and the pellet was washed with 1 mL buffer. The washing process was repeated twice, and the concentration of purified AuNP@PK was characterized to be 12.2 nM using $\varepsilon_{520} = 3.945 \times 10^8\ M^{-1}cm^{-1}$. Based on the binding capacity of PK, 67.7 µL of AuNP@PK was used for the 200 µL proteolytic reaction to ensure total PK concentration is approximately 0.01 mg/mL, the same value as for the no-NP sample and the in-situ AuNP@PK sample.

Subsequently, 200 µL of PK proteolytic assay was prepared by mixing the PK stock solutions (free PK, in-situ AuNP@PK or purified AuNP@PK) with BSA in buffer solution (10 mM $KH_2PO_4$, pH 7.5) at room temperature. 10 µL of reaction solution was sampled at an incubation time of 1 min, 2 mins, and 5 mins. The sampled reaction solution was immediately mixed with an equal amount of 2X SDS loading dye (Bio-Rad) containing 5% beta-mercaptoethanol (BME) and heated at 95°C for 10 mins to quench the reaction. Then the reaction solutions were analyzed by SDS-PAGE.

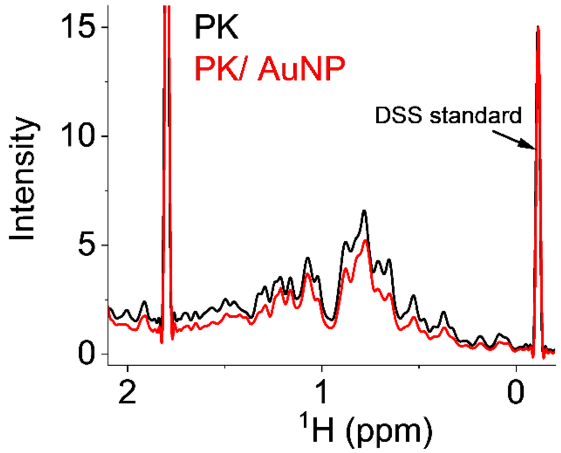

**Supplementary Fig. 4 |** 1D proton NMR spectra of 20 µM PK sample (black, PK) and 20 µM PK mixed with 50 nM AuNP (red, PK/AuNP). The signal is calibrated with DSS standard in each spectrum, and signal reduction is caused by the bound PK, with which the binding capacity $C_{PK}$ was calculated.

**Human carbonic anhydrase (HCA) activity assay**

Two equivalences of $ZnCl_2$ were added to apo-HCA for activity measurements. The HCA concentration was determined using $\varepsilon_{280}$ of $54{,}000\ M^{-1}cm^{-1}$. A 10 mM $p$NPA (substrate) stock solution was prepared by dissolving $p$NPA in acetonitrile. A 1,000 µL activity reaction was run using the following conditions: 0.2 µM HCA (free or bound) was mixed with 100 µM $p$NPA in 10 mM HEPES buffer at pH 7.5 in a disposable cuvette. Once HCA and $p$NPA were mixed, a series of UV-vis spectra (from 400 nm to 650 nm) were taken every 10 s for 10 mins. The extinction at 404 nm was monitored as a function of time to evaluate the esterase activity of HCA, as the colorless $p$NPA was converted to yellow 4-nitrophenol ($p$NP $\varepsilon_{404} = 17{,}300\ M^{-1}cm^{-1}$).

The binding capacity of HCA towards AuNPs was quantified by 1D NMR identically as for PK, and it was found to be 42 ± 8 HCA per NP (**Supplementary Fig. 5**), and the two AuNP-bound HCA samples (in-situ and purified) were prepared in the same way, except that in the in-situ method 100% HCA was bound to AuNPs.

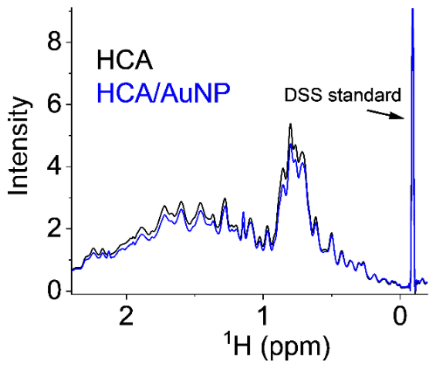

**Supplementary Fig. 5 |** 1D proton NMR spectra of 20 µM HCA sample (black, HCA) and 20 µM HCA mixed with 50 nM AuNP (red, HCA/AuNP). The signal is calibrated with DSS standard in each spectrum, and signal reduction corresponds to bound HCA concentration.

### Quantification of Charge Effect on GB3 binding onto AuNPs

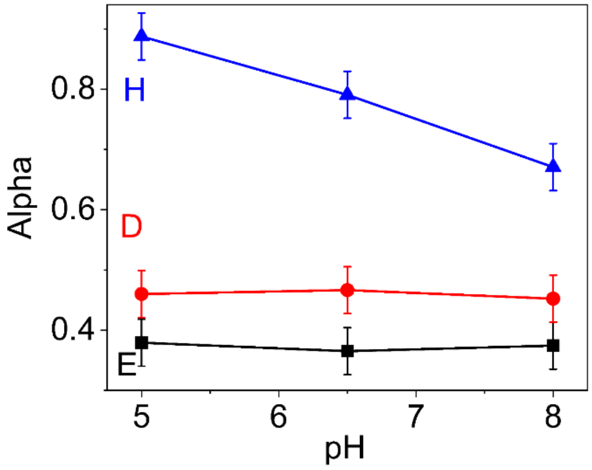

**Supplementary Fig. 6 |** Effect of pH change (from 5-8) on GB3 competitive binding with AuNPs. The blue, red and black data points represent K13H, K13D and K13E GB3 variants.

**Kinetic study of GB3 binding onto AuNP individually or in competition with SOFAST-HMQC**

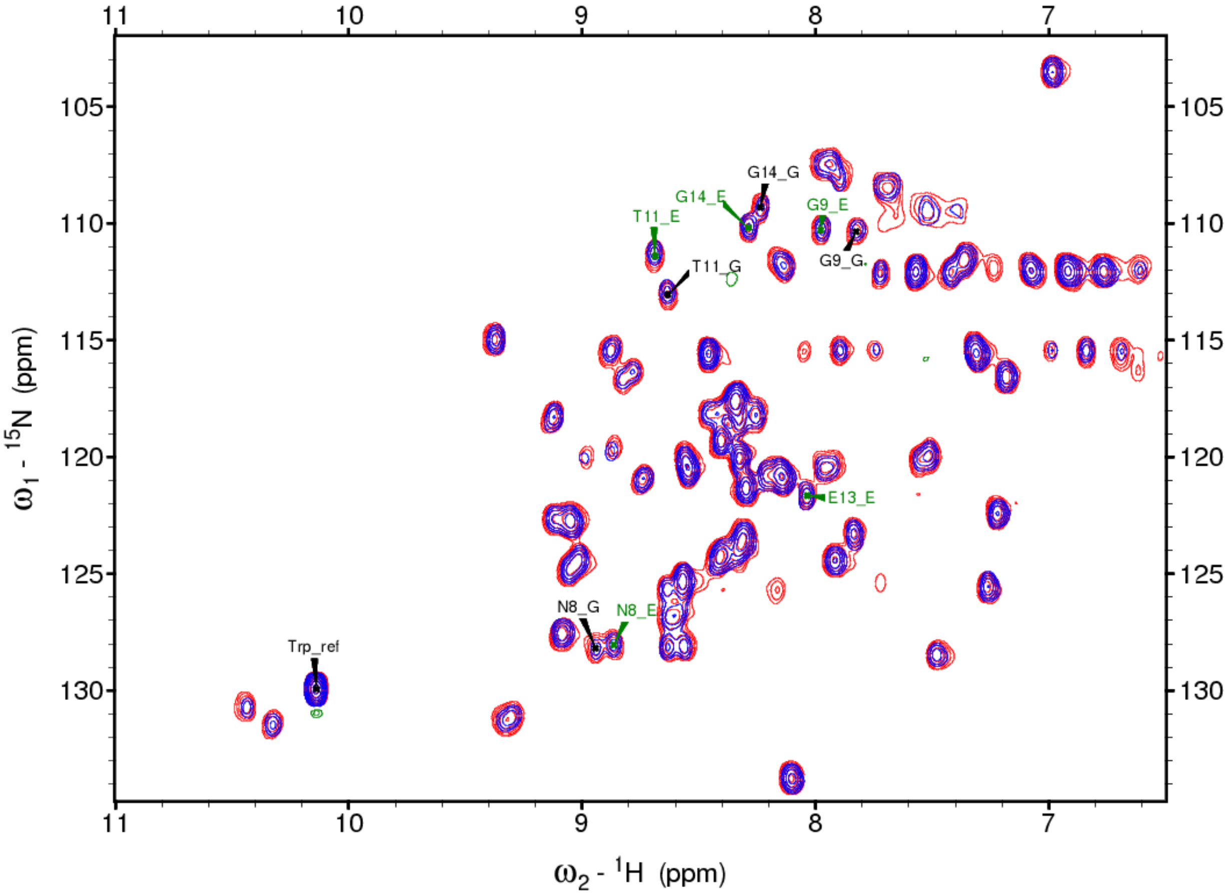

**Supplementary Fig. 7 |** Complete SOFAST-HMQC spectra of $^{15}N$ K13E/$^{15}N$ K13G mixture without (red) and with (blue) AuNPs after incubation of 365 mins. The signal reduction for the blue spectrum as compared to the red one is due to protein adsorption onto AuNPs. Well-resolved peak assignments for K13G ("_G") and K13E ("_E") are used for quantitative analysis. Because the proteins only differ by one residue, the majority of peaks overlap.

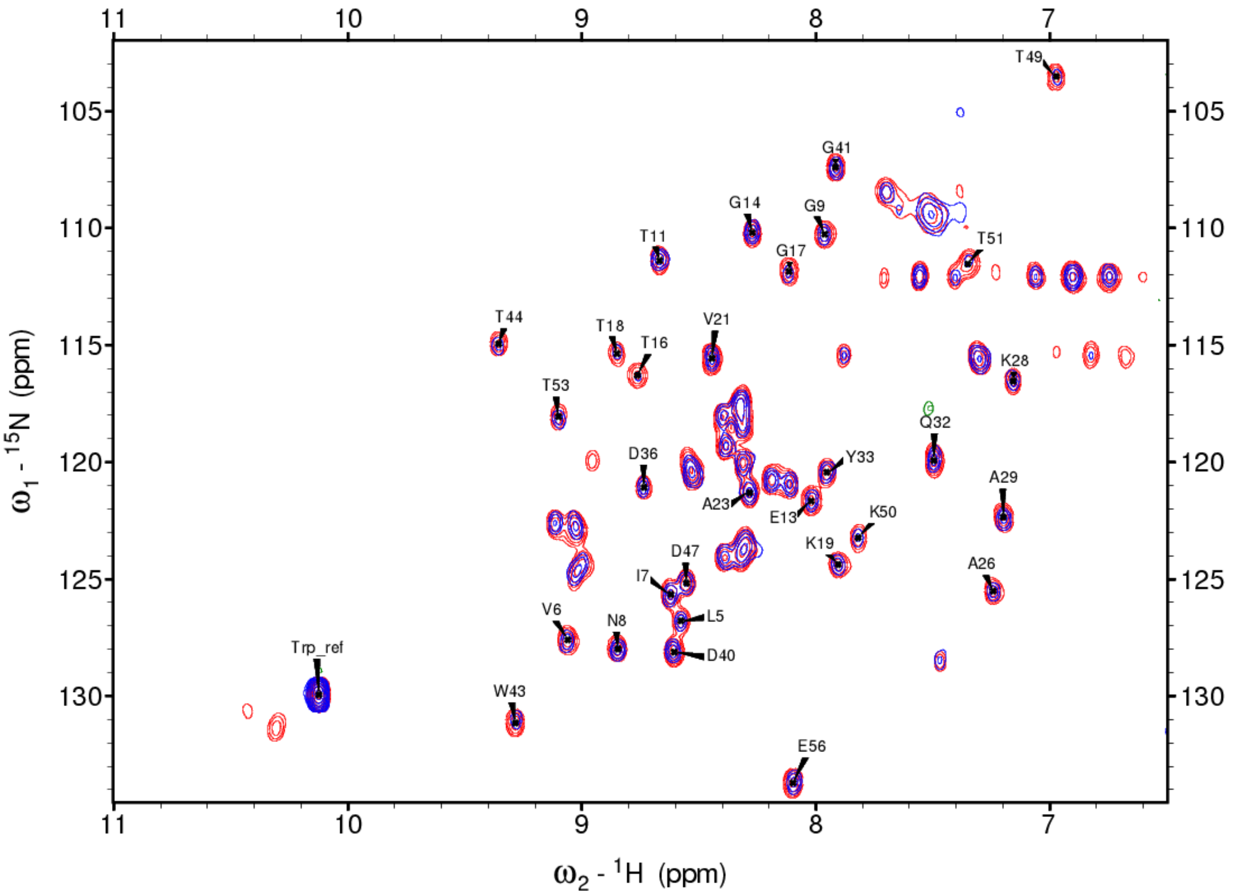

**Supplementary Fig. 8 |** Representative SOFAST-HMQC spectra of $^{15}$N K13E without (red) and with (blue) AuNPs for 365 mins. The intensity of Trp reference is used to calibrate residue peak intensities. The assigned peaks are well resolved and used for quantitative analysis.

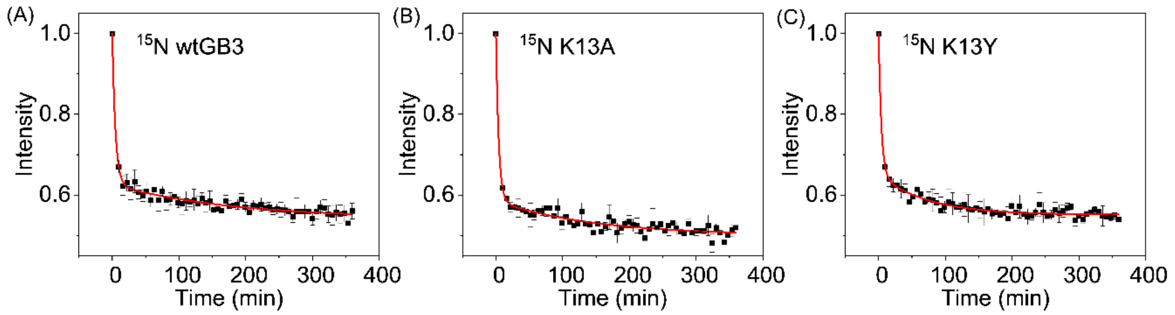

**Supplementary Fig. 9 |** Normalized average peak intensities of 20 µM of (A) $^{15}$N wt (B) $^{15}$N K13A, and (C) $^{15}$N K13Y as a function of incubation time after mixing with 50 nM AuNPs. The 0 min intensity is acquired with protein samples without AuNPs, by which all peak intensities are normalized.

**Supplementary Table 5 |** Kinetics time constants of GB3 variants using two-process decay model

| Parameter | K13Y | K13A | wtGB3 | K13G | K13E | K13E-mixture | K13G-mixture |
|---|---|---|---|---|---|---|---|
| **$y_0$** | 0.55±0.01 | 0.50±0.01 | 0.54±0.01 | 0.50±0.01 | 0.50±0.01 | 0.86±0.01 | 0.70±0.01 |
| **$A_1$** | 0.35±0.01 | 0.41±0.01 | 0.37±0.01 | 0.39±0.01 | 0.40±0.01 | 0.13±0.07 | 0.23±0.02 |
| **$t_1$ (min)** | **4.2±0.6** | **4.2±0.6** | **4.8±0.5** | **4.1±0.5** | **4.8±0.5** | **9±7** | **10±2** |
| **$A_2$** | 0.10±0.01 | 0.08±0.01 | 0.08±0.01 | 0.10±0.01 | 0.09±0.01 | 0.01±0.07 | 0.07±0.01 |
| **$t_2$ (min)** | 70±10 | 150±50 | 180±50 | 90±14 | 140±30 | 50±400 | 200±50 |
| **$R^2$** | 0.98 | 0.97 | 0.98 | 0.98 | 0.97 | 0.46 | 0.82 |

The kinetic data are fitted with equation $y = y_0 + A_1 e^{(-\frac{x}{t_1})} + A_2 e^{(-\frac{x}{t_2})}$, where $x$ refers to incubation time in mins and $y$ to normalized residue intensities. Only the pseudo first-order time ($t_1$ and $t_2$) reflect the kinetics of binding. The errors in Table 5 represent fitting errors and may not reflect experimental uncertainties from repeated measurements.

### Characterization of Purified AuNP@PK

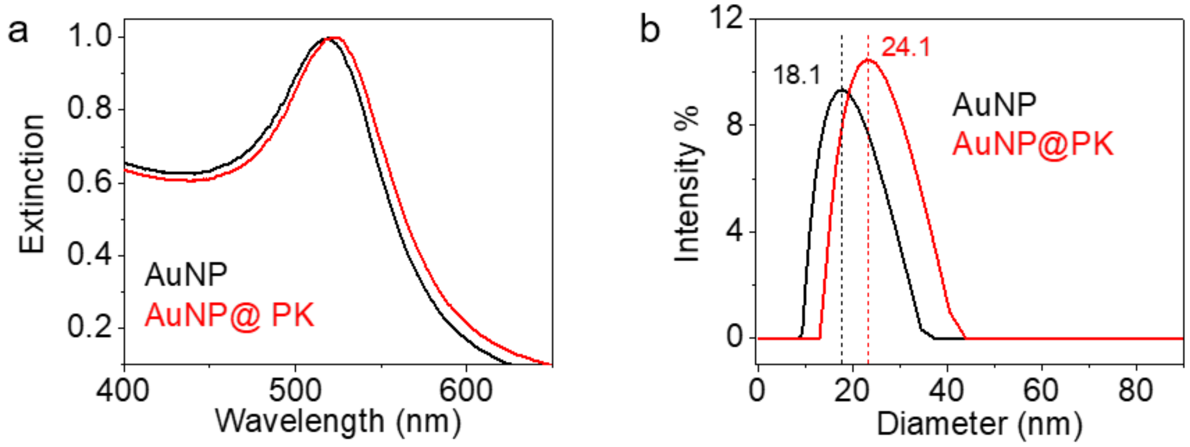

**Supplementary Fig. 10 | a,** UV-vis spectra, and **b,** DLS analysis of the purified pre-coated AuNP@PK after washing. The red shift of peak extinction wavelength of AuNP in UV-vis and increased hydrodynamic diameter of AuNP are caused by the adsorption of PK.

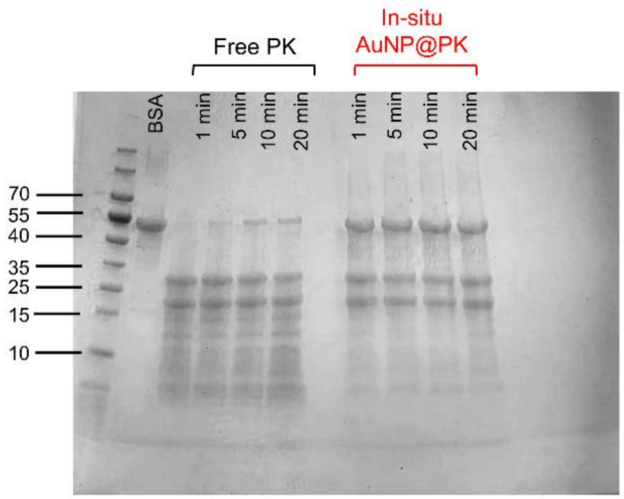

**Supplementary Fig. 11 | AuNP-bound BSA is protected from proteolysis by PK when excess AuNP is added in preparation of in-situ AuNP@PK.** With the binding capacity determined with 1D NMR, 6.8 nM AuNP is required to fully bind 0.01 mg/mL PK. Here, we incubated 0.01 mg/mL PK with 30 nM AuNP (> 4 times in excess) before adding BSA. In contrast to the results presented in the main text where AuNP is not in excess, a significant fraction of AuNP-bound BSA is not cleaved by PK, and the digestion is incomplete.

**Characterization of purified AuNP@HCA**

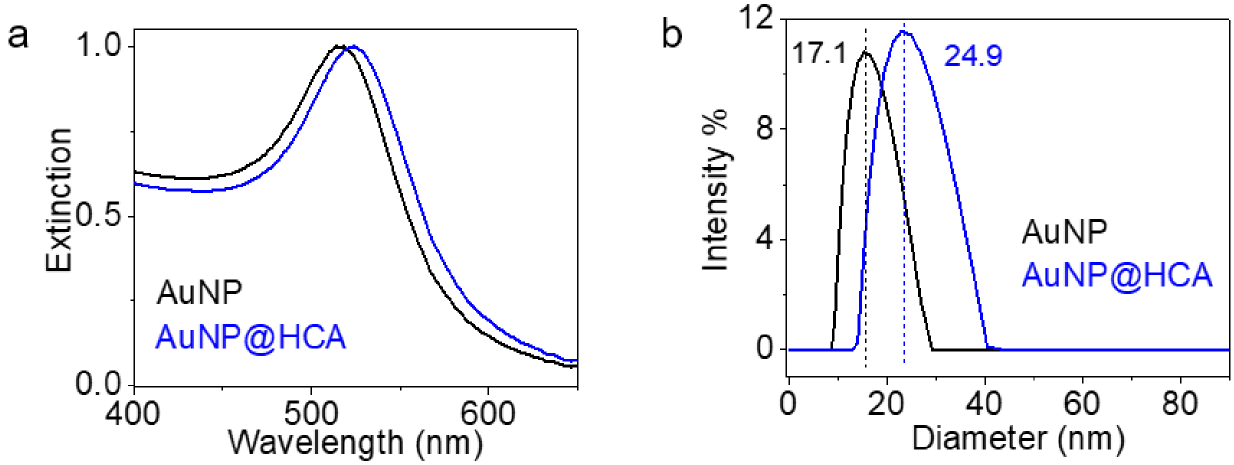

**Supplementary Fig. 12 | a,** UV-vis spectra, and **b,** DLS analysis of the purified pre-coated AuNP@HCA after washing. The red shift of peak extinction wavelength of AuNP in UV-vis and increased hydrodynamic diameter of AuNP are caused by the binding of HCA.

### Survey of Literature-Reported AuNP-bound Enzyme Activities

| **Enzyme 1** | **Alcohol dehydrogenase (TbADH)** |
|---|---|
| PDB ID | 1YKF |
| NP properties | 15-nm-Mercaptopropionic acid-capped AuNP |
| AuNP-bound activity from literature | Activity decreases from 12.5 U to 3.5U when bound.[11] |
| Active site(s) | Residues 37, 59, 150 for $Zn^{2+}$ binding, and residues 218, 340 for NADP binding.[12] |
| Analysis | Active sites should experience some steric hindrance from AuNP, but will not be completely blocked. |

### Predicted Binding Surface:

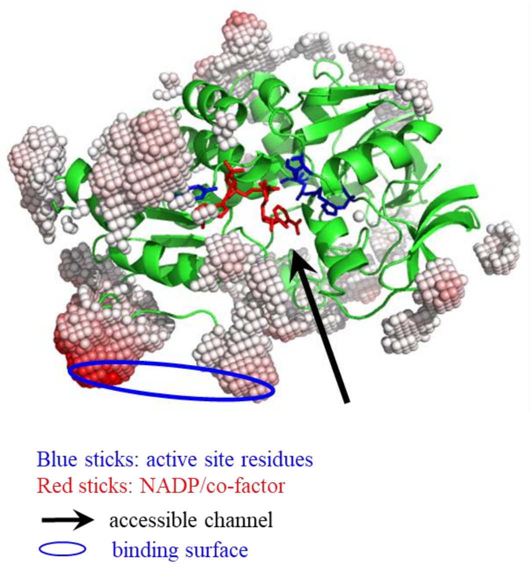

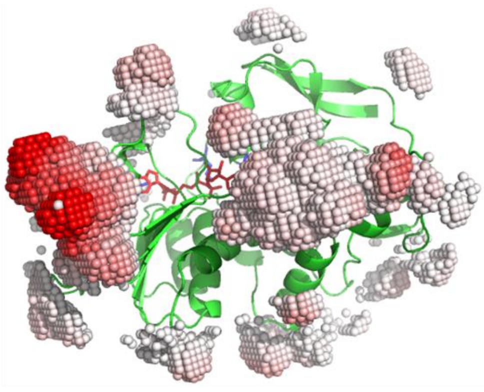

| Enzyme 2 | Acetylcholinesterase (AChE) |
|---|---|
| PDB ID | 1EEA |
| NP properties | 14 nm citrate capped AuNP |
| AuNP-bound activity from literature | Activity is fully retained.[13] |
| Active site(s) | Residues Ser 200, His 440, and Glu 327 form a catalytic triad.[14] |
| Analysis | The channel to the catalytic triad should not be blocked. |

**Predicted Binding Surface:**

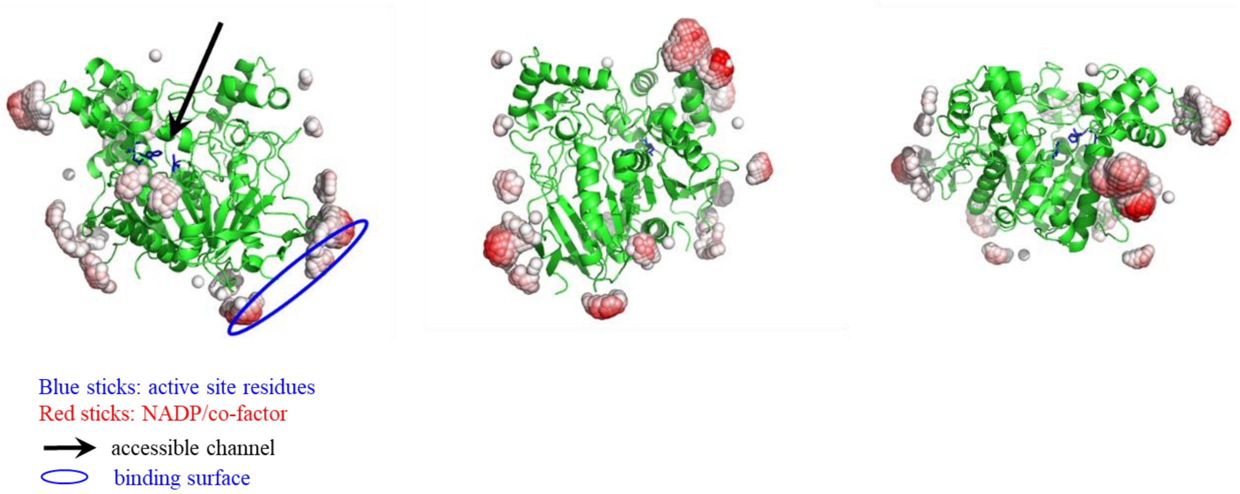

| Enzyme 3 | Citrate synthase (CS) |
|---|---|
| PDB ID | 1CTS |
| NP properties | 14 nm citrate capped AuNP |
| AuNP-bound activity from literature | Activity is fully retained.[13] |
| Active site(s) | His 274, His 320, and Asp 375[15] |
| Analysis | The channel to the catalytic triad should not be blocked. |

**Predicted Binding Surface:**

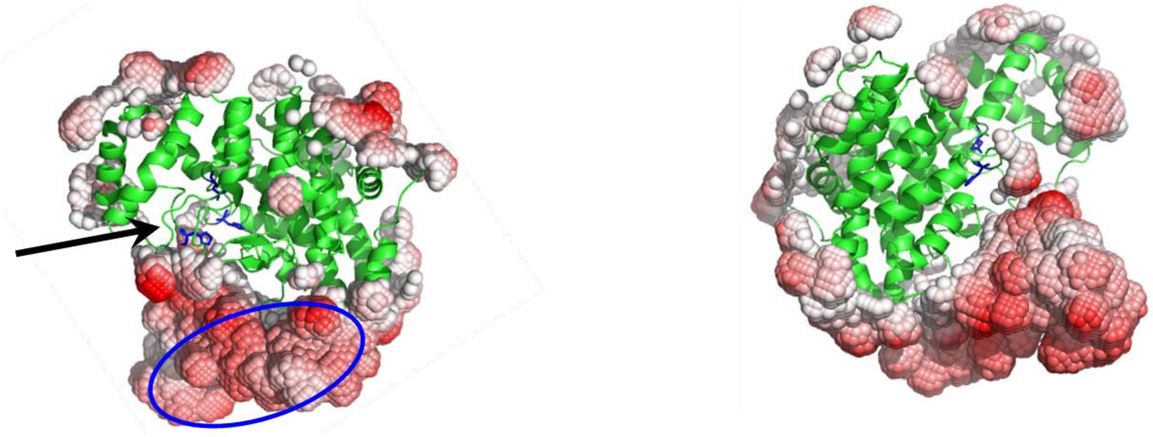

Blue sticks: active site residues
accessible channel
binding surface

| Enzyme 4 | Horseradish peroxidase (HRP) |
|---|---|
| PDB ID | 1HCH |
| NP properties | 14 nm citrate-capped AuNP |
| AuNP-bound activity from literature | Activity is only 50% retained, and further decreases to 5% when HRP packing density increases[16] |
| Active site(s) | A hydrophobic pocket formed by His 42, Phe 68, Gly 69, Ala 140, Pro 141, Phe 142, and Phe 179[17] |
| Analysis | The access to the heme and active site should be sterically hindered. Increasing packing density may enhance hindrance from nearby HRP, which decreases activity but not considered by our model. |

**Predicted Binding Surface:**

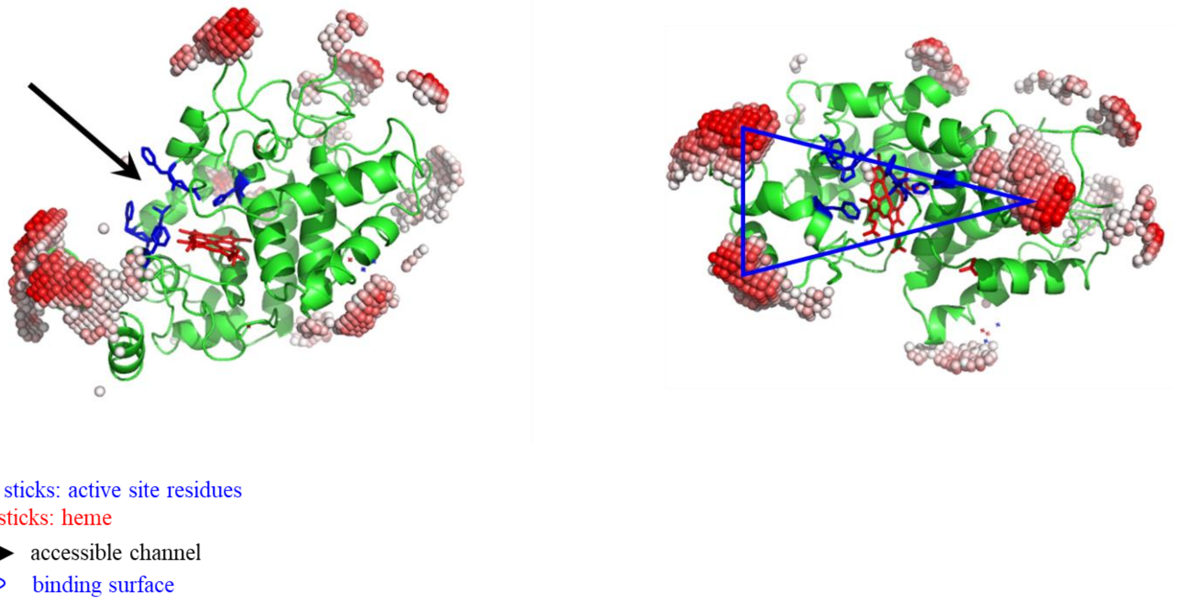

| **Enzyme 5** | **Cytochrome *c* (Cyt C)** |
|---|---|
| PDB ID | 1OCD |
| NP properties | 10-nm citrate-capped silver NP (AgNP) |
| Bound orientation from literature | The heme ring plane lies at a slight angle to the NP surface at low coverage. It re-orients to a more vertical orientation at high coverage.[18] |
| Analysis | Cyt *c* should bind at the back (blue circle), where the highest binding affinity region lies. This will orient the heme ring towards AgNP, which is consistent with observations. However, Cyt *c* has several regions of high affinity binding patches, which may explain its reorientation. |

**Predicted Binding Surface:**

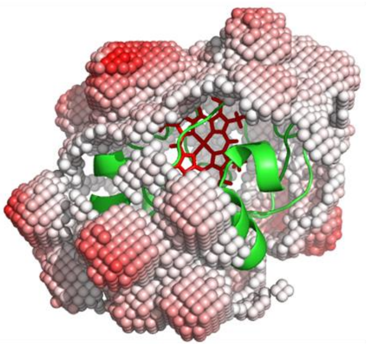

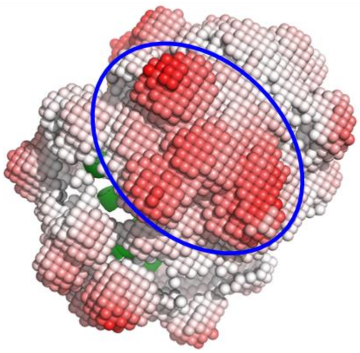

Blue sticks: active site residues
Red sticks: heme
accessible channel
binding surface

| **Enzyme 6** | **Glucose Oxidase (GOx)** |
|---|---|
| PDB ID | 1GAL |
| NP properties | 10-nm citrate-capped AuNP |
| AuNP-bound activity from literature | Activity decreases by half. Authors attributed the loss of function to protein unfolding on NP.[19] |
| Active site(s) | His 516, Glu 412, and His 559[20] |
| Analysis | Our model predicts no steric hindrance and does not explain the observed result. |

**Predicted Binding Surface:**

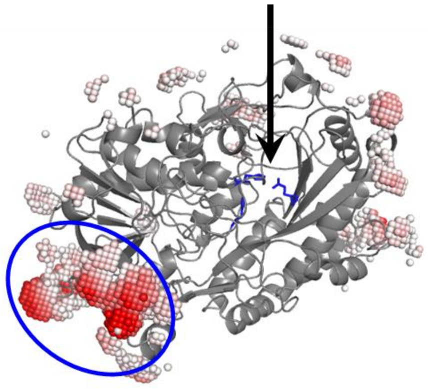

Blue sticks: active site residues

accessible channel

binding surface

$^{1}H$ and $^{15}N$ Chemical Shifts of 20 GB3 variants

**Supplementary Table 6.** Chemical Shifts of WT GB3, K13H, and K13A GB3 variants

| Residues in WT GB3 | | WT GB3 $^{15}N$ (ppm) | WT GB3 $^{1}H$ (ppm) | K13H GB3 $^{15}N$ (ppm) | K13H GB3 $^{1}H$ (ppm) | K13A GB3 $^{15}N$ (ppm) | K13A GB3 $^{1}H$ (ppm) |
|---|---|---|---|---|---|---|---|
| 2 | Gln | 123.27 | 8.31 | 123.54 | 8.35 | 124.02 | 8.39 |
| 3 | Tyr | 124.31 | 9.04 | 124.32 | 9.04 | 124.33 | 9.04 |
| 4 | Lys | 122.85 | 9.09 | 122.71 | 9.10 | 122.58 | 9.07 |
| 5 | Leu | 126.82 | 8.61 | 126.66 | 8.61 | 126.79 | 8.64 |
| 6 | Val | 127.32 | 9.15 | 127.12 | 9.10 | 127.49 | 9.11 |
| 7 | Ile | 125.73 | 8.76 | 125.50 | 8.69 | 125.61 | 8.67 |
| 8 | Asn | 129.35 | 8.98 | 129.20 | 8.92 | 128.29 | 8.91 |
| 9 | Gly | 110.52 | 7.91 | 110.29 | 7.83 | 110.15 | 7.80 |
| 10 | Lys | 120.97 | 9.52 | 120.24 | 9.36 | 120.03 | 9.12 |
| 11 | Thr | 109.12 | 8.76 | 109.16 | 8.77 | 111.51 | 8.72 |
| 12 | Leu | 125.95 | 7.57 | 124.85 | 7.47 | 125.12 | 7.76 |
| 13 | Lys | 123.84 | 8.09 | 121.60 | 8.39 | 124.12 | 7.95 |
| 14 | Gly | 109.46 | 8.30 | 110.45 | 8.19 | 108.50 | 8.24 |
| 15 | Glu | 118.33 | 8.38 | 118.28 | 8.35 | 118.56 | 8.35 |
| 16 | Thr | 115.83 | 8.81 | 115.92 | 8.77 | 116.51 | 8.84 |
| 17 | Thr | 111.99 | 8.11 | 111.96 | 8.12 | 111.65 | 8.15 |
| 18 | Thr | 115.28 | 8.91 | 115.29 | 8.92 | 115.44 | 8.91 |
| 19 | Lys | 124.53 | 7.93 | 124.49 | 7.93 | 124.48 | 7.93 |
| 20 | Ala | 124.95 | 9.08 | 124.84 | 9.09 | 124.92 | 9.09 |
| 21 | Val | 115.46 | 8.46 | 115.71 | 8.49 | 115.56 | 8.46 |
| 22 | Asp | 115.58 | 7.32 | 115.56 | 7.31 | 115.58 | 7.32 |
| 23 | Ala | 121.33 | 8.32 | 121.34 | 8.31 | 121.32 | 8.31 |
| 24 | Glu | 119.28 | 8.42 | 119.23 | 8.41 | 119.29 | 8.41 |
| 25 | Thr | 117.60 | 8.34 | 117.62 | 8.34 | 117.79 | 8.36 |
| 26 | Ala | 125.47 | 7.23 | 125.46 | 7.24 | 125.52 | 7.26 |
| 27 | Glu | 117.60 | 8.34 | 117.55 | 8.34 | 117.41 | 8.36 |
| 28 | Lys | 116.55 | 7.17 | 116.59 | 7.17 | 116.57 | 7.19 |
| 29 | Ala | 122.39 | 7.21 | 122.41 | 7.19 | 122.43 | 7.23 |
| 30 | Phe | 119.97 | 8.58 | 119.93 | 8.58 | 120.02 | 8.58 |
| 31 | Lys | 123.13 | 9.01 | 123.16 | 9.01 | 122.91 | 9.06 |
| 32 | Gln | 119.80 | 7.48 | 119.80 | 7.47 | 120.00 | 7.54 |
| 33 | Tyr | 120.60 | 8.05 | 120.61 | 8.04 | 120.47 | 7.98 |
| 34 | Ala | 122.69 | 9.18 | 122.74 | 9.20 | 122.64 | 9.13 |
| 35 | Asn | 118.18 | 8.37 | 118.19 | 8.37 | 118.11 | 8.45 |
| 36 | Asp | 121.45 | 8.82 | 121.47 | 8.79 | 121.10 | 8.79 |

| Residues in WT GB3 | | WT GB3 $^{15}N$ (ppm) | WT GB3 $^{1}H$ (ppm) | K13H GB3 $^{15}N$ (ppm) | K13H GB3 $^{1}H$ (ppm) | K13A GB3 $^{15}N$ (ppm) | K13A GB3 $^{1}H$ (ppm) |
|---|---|---|---|---|---|---|---|
| 37 | Gln | 115.53 | 7.37 | 115.58 | 7.37 | 115.23 | 7.33 |
| 38 | Gly | 108.38 | 7.78 | 108.43 | 7.78 | 108.44 | 7.72 |
| 39 | Val | 120.95 | 8.10 | 120.87 | 8.09 | 120.94 | 8.14 |
| 40 | Asp | 128.301 | 8.723 | 128.078 | 8.675 | 128.331 | 8.676 |
| 41 | Gly | 107.470 | 7.896 | 107.516 | 7.930 | 107.322 | 7.949 |
| 42 | Val | 120.624 | 8.238 | 120.714 | 8.219 | 120.607 | 8.215 |
| 43 | Trp | 131.294 | 9.308 | 131.342 | 9.323 | 131.210 | 9.314 |
| 44 | Thr | 114.763 | 9.371 | 114.909 | 9.393 | 114.911 | 9.383 |
| 45 | Tyr | 120.386 | 8.551 | 120.490 | 8.564 | 120.537 | 8.554 |
| 46 | Asp | 128.541 | 7.560 | 128.515 | 7.553 | 128.498 | 7.510 |
| 47 | Asp | 125.161 | 8.571 | 125.149 | 8.577 | 125.146 | 8.572 |
| 48 | Ala | 120.024 | 8.319 | 120.068 | 8.324 | 120.071 | 8.327 |
| 49 | Thr | 103.394 | 6.987 | 135.658 | 6.995 | 135.687 | 6.994 |
| 50 | Lys | 123.258 | 7.849 | 123.254 | 7.851 | 123.273 | 7.847 |
| 51 | Thr | 111.486 | 7.392 | 111.424 | 7.388 | 111.410 | 7.369 |
| 52 | Phe | 131.383 | 10.363 | 131.400 | 10.371 | 131.497 | 10.351 |
| 53 | Thr | 117.798 | 9.145 | 117.986 | 9.149 | 118.099 | 9.136 |
| 54 | Val | 123.362 | 8.242 | 123.680 | 8.289 | 123.380 | 8.334 |
| 55 | Thr | 124.326 | 8.325 | 124.370 | 8.342 | 124.221 | 8.353 |
| 56 | Glu | 133.835 | 7.867 | 133.952 | 7.914 | 133.926 | 8.058 |

**Supplementary Table 7.** $^{1}H$ and $^{15}N$ Chemical Shift values of K13F, K13N, and K13Y GB3 variants

| Residues in WT GB3 | | K13F GB3 $^{15}N$ (ppm) | K13F GB3 $^{1}H$ (ppm) | K13N GB3 $^{15}N$ (ppm) | K13N GB3 $^{1}H$ (ppm) | K13Y GB3 $^{15}N$ (ppm) | K13Y GB3 $^{1}H$ (ppm) |
|---|---|---|---|---|---|---|---|
| 2 | Gln | 123.29 | 8.33 | 123.34 | 8.33 | - | - |
| 3 | Tyr | 124.30 | 9.03 | 124.33 | 9.05 | 124.29 | 9.04 |
| 4 | Lys | 122.60 | 9.08 | 122.66 | 9.08 | 122.72 | 9.10 |
| 5 | Leu | 126.63 | 8.58 | 126.82 | 8.62 | 126.70 | 8.57 |
| 6 | Val | 127.29 | 9.04 | 127.40 | 9.09 | 127.11 | 9.00 |
| 7 | Ile | 125.75 | 8.65 | 125.58 | 8.66 | 125.85 | 8.67 |
| 8 | Asn | 127.76 | 8.86 | 128.44 | 8.87 | 126.92 | 8.80 |
| 9 | Gly | 110.25 | 7.91 | 110.67 | 7.98 | 113.00 | 7.88 |
| 10 | Lys | 120.21 | 9.23 | 120.51 | 9.30 | 120.87 | 9.35 |
| 11 | Thr | 109.50 | 8.76 | 110.01 | 8.80 | 108.86 | 8.80 |
| 12 | Leu | 124.80 | 7.45 | 124.72 | 7.53 | 124.31 | 7.38 |
| 13 | Lys | 122.31 | 7.98 | 121.18 | 8.21 | 123.32 | 8.05 |
| 14 | Gly | 111.31 | 8.05 | 109.31 | 8.33 | 110.47 | 7.99 |
| 15 | Glu | 118.12 | 8.23 | 118.38 | 8.36 | 117.27 | 8.13 |
| 16 | Thr | 116.25 | 8.74 | 116.15 | 8.82 | 115.94 | 8.69 |
| 17 | Thr | 112.00 | 8.12 | 111.90 | 8.12 | 111.93 | 8.11 |
| 18 | Thr | 115.36 | 8.90 | 115.37 | 8.91 | 115.31 | 8.91 |
| 19 | Lys | 124.52 | 7.92 | 124.55 | 7.93 | 124.60 | 7.93 |
| 20 | Ala | 124.90 | 9.08 | 124.93 | 9.09 | 124.94 | 9.07 |
| 21 | Val | 115.61 | 8.47 | 115.54 | 8.47 | 115.50 | 8.45 |
| 22 | Asp | 115.61 | 7.32 | 115.57 | 7.32 | 115.56 | 7.31 |
| 23 | Ala | 121.33 | 8.31 | 121.33 | 8.32 | 121.35 | 8.32 |
| 24 | Glu | 119.26 | 8.40 | 119.30 | 8.41 | 119.24 | 8.40 |
| 25 | Thr | 117.40 | 8.34 | 117.76 | 8.36 | 117.64 | 8.33 |
| 26 | Ala | 125.48 | 7.24 | 125.50 | 7.26 | 125.48 | 7.22 |
| 27 | Glu | 117.71 | 8.34 | 117.42 | 8.36 | 117.64 | 8.33 |
| 28 | Lys | 116.55 | 7.17 | 116.59 | 7.18 | 116.53 | 7.16 |
| 29 | Ala | 122.40 | 7.21 | 122.43 | 7.22 | 122.39 | 7.19 |
| 30 | Phe | 119.94 | 8.57 | 120.00 | 8.59 | 119.94 | 8.57 |
| 31 | Lys | 123.06 | 9.02 | 123.11 | 9.05 | 123.15 | 9.01 |
| 32 | Gln | 119.82 | 7.49 | 119.92 | 7.51 | 119.76 | 7.46 |
| 33 | Tyr | 120.56 | 8.02 | 120.60 | 8.03 | 120.65 | 8.03 |
| 34 | Ala | 122.71 | 9.17 | 122.73 | 9.18 | 122.72 | 9.18 |
| 35 | Asn | 118.12 | 8.38 | 118.16 | 8.40 | 118.12 | 8.34 |
| 36 | Asp | 121.39 | 8.78 | 121.34 | 8.80 | 121.46 | 8.77 |
| 37 | Gln | 115.53 | 7.36 | 115.42 | 7.36 | 115.61 | 7.38 |
| 38 | Gly | 108.47 | 7.77 | 108.57 | 7.77 | 108.59 | 7.78 |
| 39 | Val | 120.90 | 8.09 | 121.00 | 8.13 | 120.78 | 8.06 |

| Residues in WT GB3 | | K13F GB3 $^{15}N$ (ppm) | K13F GB3 $^{1}H$ (ppm) | K13N GB3 $^{15}N$ (ppm) | K13N GB3 $^{1}H$ (ppm) | K13Y GB3 $^{15}N$ (ppm) | K13Y GB3 $^{1}H$ (ppm) |
|---|---|---|---|---|---|---|---|
| 40 | Asp | 128.05 | 8.66 | 128.43 | 8.74 | 127.74 | 8.61 |
| 41 | Gly | 107.47 | 7.94 | 107.53 | 7.93 | 107.50 | 7.95 |
| 42 | Val | 120.76 | 8.21 | 120.63 | 8.22 | 120.86 | 8.17 |
| 43 | Trp | 131.26 | 9.31 | 131.13 | 9.30 | 131.38 | 9.32 |
| 44 | Thr | 114.94 | 9.39 | 114.87 | 9.39 | 115.02 | 9.42 |
| 45 | Tyr | 120.54 | 8.55 | 120.50 | 8.55 | 120.56 | 8.55 |
| 46 | Asp | 128.51 | 7.53 | 128.53 | 7.53 | 128.50 | 7.53 |
| 47 | Asp | 125.12 | 8.57 | 125.13 | 8.57 | 125.10 | 8.56 |
| 48 | Ala | 120.05 | 8.32 | 120.06 | 8.33 | 120.06 | 8.32 |
| 49 | Thr | 135.69 | 7.00 | 135.67 | 7.00 | 135.70 | 6.99 |
| 50 | Lys | 123.21 | 7.85 | 123.23 | 7.85 | 123.17 | 7.84 |
| 51 | Thr | 111.41 | 7.38 | 111.43 | 7.38 | 111.35 | 7.36 |
| 52 | Phe | 131.42 | 10.35 | 131.44 | 10.37 | 131.48 | 10.35 |
| 53 | Thr | 117.95 | 9.13 | 117.94 | 9.13 | 118.05 | 9.14 |
| 54 | Val | 123.92 | 8.34 | 124.05 | 8.31 | 124.00 | 8.39 |
| 55 | Thr | 124.21 | 8.35 | 123.87 | 8.38 | 124.24 | 8.36 |
| 56 | Glu | 134.01 | 8.00 | 133.87 | 8.08 | 134.28 | 7.93 |

**Supplementary Table 8.** $^{1}$H and $^{15}$N Chemical Shift values of K13W, K13T, and K13S GB3 variants

| Residues in WT GB3 | | K13W GB3 $^{15}$N (ppm) | K13W GB3 $^{1}$H (ppm) | K13T GB3 $^{15}$N (ppm) | K13T GB3 $^{1}$H (ppm) | K13S GB3 $^{15}$N (ppm) | K13S GB3 $^{1}$H (ppm) |
|---|---|---|---|---|---|---|---|
| 2 | Gln | 123.85 | 8.32 | 123.12 | 8.33 | 123.50 | 8.35 |
| 3 | Tyr | 124.30 | 9.03 | 124.29 | 9.04 | 124.33 | 9.04 |
| 4 | Lys | 122.60 | 9.09 | 122.76 | 9.07 | 122.67 | 9.08 |
| 5 | Leu | 126.71 | 8.59 | 126.81 | 8.61 | 126.83 | 8.63 |
| 6 | Val | 127.31 | 9.06 | 127.41 | 9.08 | 127.38 | 9.10 |
| 7 | Ile | 125.70 | 8.63 | 125.57 | 8.65 | 125.51 | 8.68 |
| 8 | Asn | 128.06 | 8.85 | 128.74 | 8.89 | 128.54 | 8.91 |
| 9 | Gly | 110.52 | 7.91 | 110.34 | 7.86 | 110.53 | 7.91 |
| 10 | Lys | 120.00 | 9.25 | 120.68 | 9.38 | 120.73 | 9.29 |
| 11 | Thr | 109.33 | 8.81 | 109.30 | 8.79 | 110.16 | 8.81 |
| 12 | Leu | 124.63 | 7.33 | 125.09 | 7.44 | 124.95 | 7.56 |
| 13 | Lys | 118.00 | 8.24 | 110.56 | 8.33 | 117.00 | 8.06 |
| 14 | Gly | 110.80 | 8.09 | 118.21 | 8.37 | 110.23 | 8.32 |
| 15 | Glu | 118.22 | 8.38 | 116.01 | 8.79 | 118.29 | 8.36 |
| 16 | Thr | 116.37 | 8.76 | 111.91 | 8.10 | 116.23 | 8.83 |
| 17 | Thr | 112.05 | 8.12 | 115.35 | 8.90 | 111.84 | 8.13 |
| 18 | Thr | 115.40 | 8.90 | 124.53 | 7.92 | 115.36 | 8.91 |
| 19 | Lys | 124.52 | 7.93 | 124.94 | 9.08 | 124.46 | 7.93 |
| 20 | Ala | 124.92 | 9.08 | 115.53 | 8.46 | 124.85 | 9.09 |
| 21 | Val | 115.56 | 8.46 | 115.50 | 7.31 | 115.66 | 8.48 |
| 22 | Asp | 115.59 | 7.32 | 121.31 | 8.31 | 115.58 | 7.32 |
| 23 | Ala | 121.33 | 8.31 | 119.30 | 8.40 | 121.32 | 8.31 |
| 24 | Glu | 119.27 | 8.40 | 117.79 | 8.34 | 119.27 | 8.41 |
| 25 | Thr | 117.81 | 8.35 | 125.48 | 7.24 | 117.65 | 8.36 |
| 26 | Ala | 125.49 | 7.24 | 116.86 | 8.02 | 125.50 | 7.26 |
| 27 | Glu | 117.45 | 8.34 | 117.46 | 8.33 | 117.55 | 8.35 |
| 28 | Lys | 116.55 | 7.17 | 116.55 | 7.17 | 116.60 | 7.18 |
| 29 | Ala | 122.40 | 7.21 | 122.41 | 7.22 | 122.42 | 7.22 |
| 30 | Phe | 119.95 | 8.58 | 120.00 | 8.58 | 119.96 | 8.59 |
| 31 | Lys | 123.04 | 9.02 | 123.11 | 9.03 | 123.04 | 9.05 |
| 32 | Gln | 119.85 | 7.49 | 119.86 | 7.50 | 119.92 | 7.52 |
| 33 | Tyr | 120.56 | 8.01 | 120.59 | 8.03 | 120.54 | 8.02 |
| 34 | Ala | 122.70 | 9.16 | 122.71 | 9.18 | 122.70 | 9.17 |
| 35 | Asn | 117.96 | 8.38 | 118.40 | 8.36 | 118.14 | 8.41 |
| 36 | Asp | 121.39 | 8.78 | 121.37 | 8.80 | 121.28 | 8.80 |
| 37 | Gln | 115.50 | 7.35 | 115.46 | 7.36 | 115.41 | 7.36 |
| 38 | Gly | 108.49 | 7.76 | 108.50 | 7.76 | 108.48 | 7.76 |
| 39 | Val | 120.96 | 8.09 | 120.99 | 8.12 | 120.98 | 8.13 |

| Residues in WT GB3 | | K13W GB3 $^{15}N$ (ppm) | K13W GB3 $^{1}H$ (ppm) | K13T GB3 $^{15}N$ (ppm) | K13T GB3 $^{1}H$ (ppm) | K13S GB3 $^{15}N$ (ppm) | K13S GB3 $^{1}H$ (ppm) |
|---|---|---|---|---|---|---|---|
| 40 | Asp | 128.19 | 8.69 | 128.37 | 8.73 | 128.27 | 8.70 |
| 41 | Gly | 107.47 | 7.92 | 107.44 | 7.89 | 107.46 | 7.93 |
| 42 | Val | 120.73 | 8.22 | 120.63 | 8.23 | 120.66 | 8.21 |
| 43 | Trp | 131.25 | 9.31 | 131.15 | 9.29 | 131.18 | 9.31 |
| 44 | Thr | 114.91 | 9.38 | 114.81 | 9.37 | 114.87 | 9.39 |
| 45 | Tyr | 120.52 | 8.55 | 120.45 | 8.54 | 120.48 | 8.56 |
| 46 | Asp | 128.51 | 7.54 | 128.56 | 7.54 | 128.51 | 7.53 |
| 47 | Asp | 125.12 | 8.57 | 125.13 | 8.56 | 125.14 | 8.58 |
| 48 | Ala | 120.06 | 8.32 | 120.04 | 8.32 | 120.08 | 8.33 |
| 49 | Thr | 135.68 | 6.99 | 135.66 | 6.99 | 135.69 | 7.00 |
| 50 | Lys | 123.23 | 7.85 | 123.22 | 7.84 | 123.24 | 7.85 |
| 51 | Thr | 111.39 | 7.37 | 111.44 | 7.38 | 111.45 | 7.38 |
| 52 | Phe | 131.47 | 10.36 | 131.41 | 10.37 | 131.44 | 10.37 |
| 53 | Thr | 117.95 | 9.13 | 117.78 | 9.12 | 117.97 | 9.14 |
| 54 | Val | 123.15 | 7.93 | 123.59 | 8.31 | 123.85 | 8.37 |
| 55 | Thr | 124.19 | 8.34 | 124.14 | 8.31 | 124.19 | 8.33 |
| 56 | Glu | 133.87 | 7.97 | 133.91 | 7.98 | 133.98 | 8.05 |

**Supplementary Table 9.** $^1$H and $^{15}$N Chemical Shift values of K13R, K13Q, and K13P GB3 variants

| Residues in WT GB3 | | K13R GB3 $^{15}$N (ppm) | K13R GB3 $^1$H (ppm) | K13Q GB3 $^{15}$N (ppm) | K13Q GB3 $^1$H (ppm) | K13P GB3 $^{15}$N (ppm) | K13P GB3 $^1$H (ppm) |
|---|---|---|---|---|---|---|---|
| 2 | Gln | 123.48 | 8.28 | 123.48 | 8.28 | 125.13 | 8.63 |
| 3 | Tyr | 124.33 | 9.04 | 124.33 | 9.04 | 124.35 | 9.02 |
| 4 | Lys | 122.72 | 9.08 | 122.72 | 9.08 | 122.09 | 9.07 |
| 5 | Leu | 126.76 | 8.62 | 126.76 | 8.62 | 126.79 | 8.74 |
| 6 | Val | 127.36 | 9.11 | 127.36 | 9.11 | 127.98 | 9.17 |
| 7 | Ile | 125.61 | 8.72 | 125.61 | 8.72 | 125.55 | 8.74 |
| 8 | Asn | 129.26 | 8.95 | 129.26 | 8.95 | 127.43 | 8.86 |
| 9 | Gly | 110.19 | 7.81 | 110.19 | 7.81 | 110.03 | 7.66 |
| 10 | Lys | 120.79 | 9.45 | 120.79 | 9.45 | 118.85 | 9.16 |
| 11 | Thr | 109.49 | 8.75 | 109.49 | 8.75 | 115.43 | 8.44 |
| 12 | Leu | 125.58 | 7.60 | 125.58 | 7.60 | 125.93 | 8.70 |
| 13 | Lys | 123.60 | 8.14 | 123.60 | 8.14 | - | - |
| 14 | Gly | 109.92 | 8.26 | 109.92 | 8.26 | 107.97 | 8.24 |
| 15 | Glu | 118.26 | 8.37 | 118.26 | 8.37 | 118.96 | 8.30 |
| 16 | Thr | 115.93 | 8.80 | 115.93 | 8.80 | 118.05 | 8.61 |
| 17 | Thr | 111.97 | 8.11 | 111.97 | 8.11 | 111.08 | 8.27 |
| 18 | Thr | 115.34 | 8.91 | 115.34 | 8.91 | 115.75 | 8.92 |
| 19 | Lys | 124.53 | 7.92 | 124.53 | 7.92 | 124.48 | 7.93 |
| 20 | Ala | 124.93 | 9.09 | 124.93 | 9.09 | 124.99 | 9.09 |
| 21 | Val | 115.55 | 8.46 | 115.55 | 8.46 | 117.26 | 8.48 |
| 22 | Asp | 115.56 | 7.31 | 115.56 | 7.31 | 114.55 | 7.24 |
| 23 | Ala | 121.32 | 8.31 | 121.32 | 8.31 | 121.31 | 8.32 |
| 24 | Glu | 119.29 | 8.41 | 119.29 | 8.41 | 119.33 | 8.41 |
| 25 | Thr | 117.62 | 8.34 | 117.62 | 8.34 | 117.69 | 8.41 |
| 26 | Ala | 125.49 | 7.24 | 125.49 | 7.24 | 125.59 | 7.32 |
| 27 | Glu | 117.62 | 8.34 | 117.62 | 8.34 | 117.57 | 8.38 |
| 28 | Lys | 116.58 | 7.17 | 116.58 | 7.17 | 116.55 | 7.25 |
| 29 | Ala | 122.41 | 7.21 | 122.41 | 7.21 | 122.46 | 7.28 |
| 30 | Phe | 119.98 | 8.58 | 119.98 | 8.58 | 120.07 | 8.59 |
| 31 | Lys | 123.10 | 9.02 | 123.10 | 9.02 | 122.40 | 9.03 |
| 32 | Gln | 119.89 | 7.50 | 119.89 | 7.50 | 120.36 | 7.66 |
| 33 | Tyr | 120.61 | 8.04 | 120.61 | 8.04 | 120.18 | 7.85 |
| 34 | Ala | 122.71 | 9.18 | 122.71 | 9.18 | 122.40 | 9.15 |
| 35 | Asn | 118.26 | 8.37 | 118.26 | 8.37 | 118.29 | 8.55 |
| 36 | Asp | 121.39 | 8.80 | 121.39 | 8.80 | 120.33 | 8.75 |
| 37 | Gln | 115.47 | 7.36 | 115.47 | 7.36 | 115.60 | 7.32 |
| 38 | Gly | 108.48 | 7.77 | 108.48 | 7.77 | 108.26 | 7.60 |
| 39 | Val | 121.00 | 8.11 | 121.00 | 8.11 | 120.34 | 8.19 |

| Residues in WT GB3 | | K13R GB3 $^{15}N$ (ppm) | K13R GB3 $^{1}H$ (ppm) | K13Q GB3 $^{15}N$ (ppm) | K13Q GB3 $^{1}H$ (ppm) | K13P GB3 $^{15}N$ (ppm) | K13P GB3 $^{1}H$ (ppm) |
|---|---|---|---|---|---|---|---|
| 40 | Asp | 128.37 | 8.73 | 128.37 | 8.73 | 128.72 | 8.63 |
| 41 | Gly | 107.51 | 7.91 | 107.51 | 7.91 | 106.88 | 8.06 |
| 42 | Val | 120.54 | 8.21 | 120.54 | 8.21 | 120.98 | 8.22 |
| 43 | Trp | 131.22 | 9.31 | 131.22 | 9.31 | 131.31 | 9.35 |
| 44 | Thr | 114.76 | 9.37 | 114.76 | 9.37 | 115.14 | 9.42 |
| 45 | Tyr | 120.41 | 8.55 | 120.41 | 8.55 | 120.76 | 8.57 |
| 46 | Asp | 128.53 | 7.55 | 128.53 | 7.55 | 128.44 | 7.45 |
| 47 | Asp | 125.15 | 8.57 | 125.15 | 8.57 | 125.15 | 8.59 |
| 48 | Ala | 120.05 | 8.32 | 120.05 | 8.32 | 120.11 | 8.34 |
| 49 | Thr | 103.39 | 6.99 | 103.39 | 6.99 | 103.47 | 7.00 |
| 50 | Lys | 123.27 | 7.84 | 123.27 | 7.84 | 123.40 | 7.85 |
| 51 | Thr | 111.44 | 7.38 | 111.44 | 7.38 | 111.29 | 7.34 |
| 52 | Phe | 131.42 | 10.37 | 131.42 | 10.37 | 131.81 | 10.34 |
| 53 | Thr | 117.81 | 9.14 | 117.81 | 9.14 | 117.94 | 8.99 |
| 54 | Val | 123.48 | 8.28 | 123.48 | 8.28 | 123.12 | 8.32 |
| 55 | Thr | 124.35 | 8.34 | 124.35 | 8.34 | 124.45 | 8.46 |
| 56 | Glu | 133.92 | 7.88 | 133.92 | 7.88 | 134.17 | 8.24 |

**Supplementary Table 10.** $^{1}$H and $^{15}$N Chemical Shift values of K13L, K13G, and K13E GB3 variants

| Residues in WT GB3 | | K13L GB3 $^{15}$N (ppm) | K13L GB3 $^{1}$H (ppm) | K13G GB3 $^{15}$N (ppm) | K13G GB3 $^{1}$H (ppm) | K13E GB3 $^{15}$N (ppm) | K13E GB3 $^{1}$H (ppm) |
|---|---|---|---|---|---|---|---|
| 2 | Gln | 123.98 | 8.35 | 123.83 | 8.35 | 123.38 | 8.34 |
| 3 | Tyr | 124.29 | 9.04 | 124.05 | 9.04 | 124.27 | 9.03 |
| 4 | Lys | 122.60 | 9.07 | 122.19 | 9.06 | 122.58 | 9.06 |
| 5 | Leu | 126.66 | 8.57 | 126.46 | 8.62 | 126.75 | 8.60 |
| 6 | Val | 127.41 | 9.08 | 127.25 | 9.09 | 127.50 | 9.09 |
| 7 | Ile | 125.76 | 8.65 | 125.50 | 8.54 | 125.60 | 8.65 |
| 8 | Asn | 128.58 | 8.89 | 128.03 | 8.94 | 128.28 | 8.88 |
| 9 | Gly | 110.44 | 7.95 | 109.87 | 7.81 | 110.20 | 7.96 |
| 10 | Lys | 120.43 | 9.20 | 120.16 | 8.99 | 120.27 | 9.12 |
| 11 | Thr | 110.09 | 8.68 | 111.98 | 8.64 | 110.74 | 8.73 |
| 12 | Leu | 125.13 | 7.57 | 124.90 | 7.95 | 125.00 | 7.59 |
| 13 | Lys | 118.73 | 8.33 | 108.73 | 7.96 | 121.94 | 8.07 |
| 14 | Gly | 109.84 | 8.37 | 108.82 | 8.19 | 109.91 | 8.29 |
| 15 | Glu | 118.03 | 8.38 | 117.86 | 8.43 | 118.45 | 8.34 |
| 16 | Thr | 116.04 | 8.72 | 116.37 | 8.79 | 116.22 | 8.78 |
| 17 | Thr | 112.15 | 8.12 | 111.58 | 8.15 | 111.99 | 8.13 |
| 18 | Thr | 115.35 | 8.89 | 115.36 | 8.90 | 115.37 | 8.88 |
| 19 | Lys | 124.59 | 7.93 | 124.40 | 7.93 | 124.50 | 7.93 |
| 20 | Ala | 124.94 | 9.08 | 124.70 | 9.08 | 124.90 | 9.07 |
| 21 | Val | 115.53 | 8.46 | 115.33 | 8.45 | 115.58 | 8.46 |
| 22 | Asp | 115.64 | 7.32 | 115.31 | 7.30 | 115.66 | 7.33 |
| 23 | Ala | 121.32 | 8.32 | 121.02 | 8.31 | 121.33 | 8.32 |
| 24 | Glu | 119.29 | 8.41 | 119.07 | 8.39 | 119.28 | 8.41 |
| 25 | Thr | 117.83 | 8.34 | 117.25 | 8.34 | 117.86 | 8.35 |
| 26 | Ala | 125.48 | 7.24 | 125.22 | 7.27 | 125.50 | 7.26 |
| 27 | Glu | 117.44 | 8.34 | 117.68 | 8.35 | 117.42 | 8.35 |
| 28 | Lys | 116.55 | 7.17 | 116.33 | 7.19 | 116.57 | 7.18 |
| 29 | Ala | 122.41 | 7.22 | 122.15 | 7.23 | 122.42 | 7.22 |
| 30 | Phe | 119.92 | 8.58 | 119.76 | 8.57 | 119.98 | 8.58 |
| 31 | Lys | 123.11 | 9.03 | 122.58 | 9.05 | 123.01 | 9.04 |
| 32 | Gln | 119.86 | 7.50 | 119.91 | 7.55 | 119.94 | 7.52 |
| 33 | Tyr | 120.59 | 8.03 | 120.26 | 7.96 | 120.54 | 8.01 |
| 34 | Ala | 122.71 | 9.19 | 122.39 | 9.13 | 122.68 | 9.16 |
| 35 | Asn | 118.23 | 8.38 | 118.13 | 8.24 | 118.12 | 8.41 |
| 36 | Asp | 121.30 | 8.78 | 120.67 | 8.74 | 121.21 | 8.78 |
| 37 | Gln | 115.50 | 7.38 | 114.97 | 7.35 | 115.39 | 7.35 |
| 38 | Gly | 108.54 | 7.77 | 108.40 | 7.70 | 108.50 | 7.75 |
| 39 | Val | 120.99 | 8.13 | 120.88 | 8.16 | 121.00 | 8.13 |

| Residues in WT GB3 | | K13L GB3 $^{15}N$ (ppm) | K13L GB3 $^{1}H$ (ppm) | K13G GB3 $^{15}N$ (ppm) | K13G GB3 $^{1}H$ (ppm) | K13E GB3 $^{15}N$ (ppm) | K13E GB3 $^{1}H$ (ppm) |
|---|---|---|---|---|---|---|---|
| 40 | Asp | 128.18 | 8.70 | 128.04 | 8.65 | 128.36 | 8.71 |
| 41 | Gly | 107.49 | 7.93 | 107.08 | 7.93 | 107.45 | 7.92 |
| 42 | Val | 120.69 | 8.22 | 120.38 | 8.20 | 120.70 | 8.23 |
| 43 | Trp | 131.23 | 9.32 | 130.78 | 9.32 | 131.09 | 9.30 |
| 44 | Thr | 114.95 | 9.39 | 114.75 | 9.38 | 114.93 | 9.37 |
| 45 | Tyr | 120.54 | 8.56 | 120.35 | 8.55 | 120.56 | 8.55 |
| 46 | Asp | 128.50 | 7.53 | 128.29 | 7.50 | 128.50 | 7.52 |
| 47 | Asp | 125.13 | 8.57 | 124.89 | 8.56 | 125.12 | 8.57 |
| 48 | Ala | 120.04 | 8.32 | 119.82 | 8.32 | 120.06 | 8.33 |
| 49 | Thr | 135.66 | 6.99 | 135.64 | 7.01 | 135.70 | 6.99 |
| 50 | Lys | 123.21 | 7.85 | 123.22 | 7.84 | 123.20 | 7.85 |
| 51 | Thr | 111.44 | 7.38 | 111.24 | 7.36 | 111.45 | 7.37 |
| 52 | Phe | 131.41 | 10.35 | 131.22 | 10.35 | 131.39 | 10.34 |
| 53 | Thr | 118.04 | 9.14 | 117.91 | 9.13 | 117.99 | 9.13 |
| 54 | Val | 125.09 | 8.00 | 123.01 | 8.32 | 124.02 | 8.33 |
| 55 | Thr | 124.26 | 8.37 | 124.08 | 8.41 | 123.94 | 8.38 |
| 56 | Glu | 133.97 | 8.04 | 133.63 | 8.11 | 133.80 | 8.10 |

**Supplementary Table 11.** $^{1}H$ and $^{15}N$ Chemical Shift values of K13C, K13D, and K13I GB3 variants

| Residues in WT GB3 | | K13C GB3 $^{15}N$ (ppm) | K13C GB3 $^{1}H$ (ppm) | K13D GB3 $^{15}N$ (ppm) | K13D GB3 $^{1}H$ (ppm) | K13I GB3 $^{15}N$ (ppm) | K13I GB3 $^{1}H$ (ppm) |
|---|---|---|---|---|---|---|---|
| 2 | Gln | 123.07 | 8.28 | 124.00 | 8.34 | 124.04 | 8.29 |
| 3 | Tyr | 124.32 | 9.03 | 124.21 | 9.02 | 124.31 | 9.04 |
| 4 | Lys | 122.72 | 9.07 | 122.50 | 9.05 | 122.81 | 9.06 |
| 5 | Leu | 126.87 | 8.61 | 126.68 | 8.56 | 126.87 | 8.59 |
| 6 | Val | 127.37 | 9.08 | 127.48 | 9.07 | 127.53 | 9.09 |
| 7 | Ile | 125.54 | 8.68 | 125.68 | 8.57 | 125.79 | 8.66 |
| 8 | Asn | 128.69 | 8.90 | 127.84 | 8.85 | 128.41 | 8.86 |
| 9 | Gly | 110.97 | 7.84 | 110.53 | 8.13 | 110.61 | 7.92 |
| 10 | Lys | 120.40 | 9.29 | 120.18 | 9.14 | 119.88 | 9.28 |
| 11 | Thr | 110.04 | 8.78 | 110.81 | 8.73 | 109.29 | 8.71 |
| 12 | Leu | 124.81 | 7.51 | 124.64 | 7.57 | 125.69 | 7.50 |
| 13 | Lys | 121.10 | 8.18 | 122.26 | 8.06 | 123.97 | 7.97 |
| 14 | Gly | 110.93 | 8.37 | 108.23 | 8.32 | 112.45 | 8.47 |
| 15 | Glu | 118.37 | 8.37 | 118.52 | 8.30 | 118.67 | 8.37 |
| 16 | Thr | 116.07 | 8.80 | 116.18 | 8.72 | 115.89 | 8.78 |
| 17 | Thr | 111.90 | 8.11 | 112.19 | 8.12 | 112.05 | 8.10 |
| 18 | Thr | 115.34 | 8.90 | 115.37 | 8.86 | 115.28 | 8.89 |
| 19 | Lys | 124.59 | 7.92 | 124.63 | 7.94 | 124.60 | 7.93 |
| 20 | Ala | 125.03 | 9.07 | 125.06 | 9.05 | 125.04 | 9.07 |
| 21 | Val | 115.30 | 8.43 | 115.28 | 8.42 | 115.28 | 8.43 |
| 22 | Asp | 115.56 | 7.31 | 115.70 | 7.33 | 115.59 | 7.31 |
| 23 | Ala | 121.33 | 8.32 | 121.35 | 8.33 | 121.33 | 8.32 |
| 24 | Glu | 119.31 | 8.41 | 119.30 | 8.40 | 119.31 | 8.41 |
| 25 | Thr | 117.57 | 8.34 | 117.69 | 8.34 | 117.68 | 8.34 |
| 26 | Ala | 125.50 | 7.24 | 125.51 | 7.26 | 125.49 | 7.23 |
| 27 | Glu | 117.57 | 8.34 | 117.48 | 8.34 | 117.68 | 8.34 |
| 28 | Lys | 116.57 | 7.17 | 116.57 | 7.18 | 116.57 | 7.17 |
| 29 | Ala | 122.42 | 7.21 | 122.44 | 7.23 | 122.41 | 7.21 |
| 30 | Phe | 120.00 | 8.58 | 119.96 | 8.57 | 119.99 | 8.57 |
| 31 | Lys | 123.12 | 9.04 | 123.11 | 9.04 | 123.19 | 9.03 |
| 32 | Gln | 119.88 | 7.50 | 119.98 | 7.52 | 119.85 | 7.49 |
| 33 | Tyr | 120.58 | 8.02 | 120.55 | 8.01 | 120.41 | 7.96 |
| 34 | Ala | 122.71 | 9.16 | 122.75 | 9.17 | 122.69 | 9.17 |
| 35 | Asn | 118.15 | 8.38 | 118.12 | 8.38 | 118.16 | 8.36 |
| 36 | Asp | 121.34 | 8.80 | 121.17 | 8.75 | 121.37 | 8.81 |
| 37 | Gln | 115.42 | 7.35 | 115.42 | 7.38 | 115.47 | 7.36 |
| 38 | Gly | 108.36 | 7.75 | 108.55 | 7.75 | 108.54 | 7.76 |
| 39 | Val | 120.99 | 8.11 | 121.04 | 8.15 | 120.64 | 8.05 |

| | | | | | | | |
|---|---|---|---|---|---|---|---|
| 40 | Asp | 128.41 | 8.74 | 128.34 | 8.71 | 128.48 | 8.76 |
| 41 | Gly | 107.45 | 7.90 | 107.52 | 7.93 | 107.52 | 7.88 |
| 42 | Val | 120.64 | 8.22 | 120.72 | 8.22 | 120.61 | 8.22 |
| 43 | Trp | 131.13 | 9.29 | 131.04 | 9.29 | 131.07 | 9.29 |
| 44 | Thr | 114.82 | 9.37 | 115.00 | 9.37 | 114.76 | 9.36 |
| 45 | Tyr | 120.47 | 8.54 | 120.67 | 8.55 | 120.41 | 8.54 |
| 46 | Asp | 128.52 | 7.53 | 128.48 | 7.50 | 128.53 | 7.54 |
| 47 | Asp | 125.14 | 8.56 | 125.11 | 8.56 | 125.14 | 8.56 |
| 48 | Ala | 120.04 | 8.32 | 120.04 | 8.32 | 120.02 | 8.31 |
| 49 | Thr | 135.67 | 6.99 | 135.70 | 6.99 | 135.65 | 6.99 |
| 50 | Lys | 123.24 | 7.84 | 123.20 | 7.84 | 123.22 | 7.84 |
| 51 | Thr | 111.45 | 7.37 | 111.41 | 7.36 | 111.49 | 7.38 |
| 52 | Phe | 131.41 | 10.35 | 131.37 | 10.31 | 131.37 | 10.35 |
| 53 | Thr | 117.87 | 9.12 | 118.13 | 9.12 | 117.77 | 9.12 |
| 54 | Val | 123.77 | 8.36 | 124.00 | 8.34 | 121.05 | 8.12 |
| 55 | Thr | 124.09 | 8.30 | 124.14 | 8.39 | 123.67 | 8.35 |
| 56 | Glu | 133.90 | 8.05 | 133.74 | 8.15 | 133.82 | 8.00 |

**Supplementary Table 12.** $^{1}H$ and $^{15}N$ Chemical Shift values of K13V and K13M GB3 variants

| Residues in WT GB3 | | K13V GB3 $^{15}N$ (ppm) | K13V GB3 $^{1}H$ (ppm) | K13M GB3 $^{15}N$ (ppm) | K13M GB3 $^{1}H$ (ppm) |
|---|---|---|---|---|---|
| 2 | Gln | 124.06 | 8.28 | 123.36 | 8.30 |
| 3 | Tyr | 124.31 | 9.04 | 124.36 | 9.03 |
| 4 | Lys | 122.80 | 9.06 | 122.81 | 9.06 |
| 5 | Leu | 126.88 | 8.59 | 126.89 | 8.60 |
| 6 | Val | 127.52 | 9.09 | 127.46 | 9.09 |
| 7 | Ile | 125.78 | 8.67 | 125.71 | 8.68 |
| 8 | Asn | 128.33 | 8.87 | 128.82 | 8.90 |
| 9 | Gly | 110.66 | 7.93 | 110.57 | 7.84 |
| 10 | Lys | 120.01 | 9.31 | 119.90 | 9.20 |
| 11 | Thr | 109.34 | 8.72 | 109.90 | 8.72 |
| 12 | Leu | 125.60 | 7.47 | 125.23 | 7.57 |
| 13 | Lys | 122.89 | 7.96 | 122.68 | 8.09 |
| 14 | Gly | 112.12 | 8.45 | 110.05 | 8.35 |
| 15 | Glu | 118.62 | 8.38 | 118.40 | 8.35 |
| 16 | Thr | 115.84 | 8.79 | 116.06 | 8.79 |
| 17 | Thr | 112.00 | 8.09 | 111.94 | 8.11 |
| 18 | Thr | 115.29 | 8.89 | 115.31 | 8.89 |
| 19 | Lys | 124.59 | 7.92 | 124.45 | 7.92 |
| 20 | Ala | 125.04 | 9.08 | 124.90 | 9.07 |
| 21 | Val | 115.27 | 8.43 | 115.55 | 8.47 |
| 22 | Asp | 115.57 | 7.32 | 115.64 | 7.31 |
| 23 | Ala | 121.31 | 8.32 | 121.32 | 8.30 |
| 24 | Glu | 119.30 | 8.41 | 119.30 | 8.41 |
| 25 | Thr | 117.65 | 8.34 | 117.46 | 8.33 |
| 26 | Ala | 125.50 | 7.23 | 125.49 | 7.24 |
| 27 | Glu | 117.65 | 8.34 | 117.69 | 8.35 |
| 28 | Lys | 116.56 | 7.17 | 116.54 | 7.16 |
| 29 | Ala | 122.40 | 7.21 | 122.39 | 7.21 |
| 30 | Phe | 120.01 | 8.57 | 119.99 | 8.57 |
| 31 | Lys | 123.17 | 9.03 | 123.10 | 9.02 |
| 32 | Gln | 119.84 | 7.49 | 119.85 | 7.49 |
| 33 | Tyr | 120.63 | 8.05 | 120.54 | 8.02 |
| 34 | Ala | 122.68 | 9.17 | 122.70 | 9.16 |
| 35 | Asn | 118.14 | 8.36 | 118.15 | 8.38 |
| 36 | Asp | 121.37 | 8.81 | 121.28 | 8.79 |
| 37 | Gln | 115.45 | 7.36 | 115.44 | 7.35 |
| 38 | Gly | 108.52 | 7.76 | 108.49 | 7.75 |
| 39 | Val | 121.04 | 8.12 | 120.97 | 8.12 |
| 40 | Asp | 128.53 | 8.78 | 128.26 | 8.70 |

| Residues in WT GB3 | | K13V GB3 $^{15}N$ (ppm) | K13V GB3 $^{1}H$ (ppm) | K13M GB3 $^{15}N$ (ppm) | K13M GB3 $^{1}H$ (ppm) |
|---|---|---|---|---|---|
| 41 | Gly | 107.49 | 7.87 | 107.47 | 7.91 |
| 42 | Val | 120.60 | 8.22 | 120.65 | 8.21 |
| 43 | Trp | 131.06 | 9.29 | 131.17 | 9.30 |
| 44 | Thr | 114.75 | 9.37 | 114.82 | 9.37 |
| 45 | Tyr | 120.39 | 8.54 | 120.43 | 8.54 |
| 46 | Asp | 128.53 | 7.54 | 128.49 | 7.52 |
| 47 | Asp | 125.13 | 8.56 | 125.16 | 8.57 |
| 48 | Ala | 120.01 | 8.31 | 120.04 | 8.32 |
| 49 | Thr | 135.64 | 6.98 | 103.45 | 6.99 |
| 50 | Lys | 123.21 | 7.84 | 123.23 | 7.84 |
| 51 | Thr | 111.48 | 7.38 | 111.54 | 7.38 |
| 52 | Phe | 131.38 | 10.36 | 131.39 | 10.35 |
| 53 | Thr | 117.72 | 9.12 | 117.90 | 9.12 |
| 54 | Val | 123.61 | 8.34 | 123.75 | 8.34 |
| 55 | Thr | 124.00 | 8.29 | 124.19 | 8.32 |
| 56 | Glu | 133.78 | 7.99 | 133.96 | 8.02 |

### References

1. Song H., Wilson D. L., Farquhar E. R., Lewis E. A., Emerson J. P. Revisiting zinc coordination in human carbonic anhydrase II. *Inorg. Chem.* **51**, 11098-11105 (2012).
2. Frens G. Controlled nucleation for the regulation of the particle size in monodisperse gold suspensions. *Nat. Phys. Sci.* **241**, 20-22 (1973).
3. Turkevich J., Stevenson P. C., Hillier J. A study of the nucleation and growth processes in the synthesis of colloidal gold. *Discuss. Faraday Soc.* **11**, 55-75 (1951).
4. Malmodin D., Papavoine C. H. M., Billeter M. Fully automated sequence-specific resonance assignments of hetero- nuclear protein spectra. *J. Biomol. NMR* **27**, 69-79 (2003).
5. Pace C. N., Vajdos F., Fee L., Grimsley G., Gray T. How to measure and predict the molar absorption coefficient of a protein. *Protein Sci.* **4**, 2411-2423 (1995).
6. Wang A., Vangala K., Vo T., Zhang D., Fitzkee N. C. A three-step model for protein–gold nanoparticle adsorption. *J. Phys. Chem. C* **118**, 8134-8142 (2014).
7. Schanda P., Kupce E., Brutscher B. SOFAST-HMQC experiments for recording two-dimensional heteronuclear correlation spectra of proteins within a few seconds. *J. Biomol. NMR* **33**, 199-211 (2005).
8. Schanda P., Van Melckebeke H., Brutscher B. Speeding up three-dimensional protein NMR experiments to a few minutes. *J. Am. Chem. Soc.* **128**, 9042-9043 (2006).
9. Xu J. X., Alom M. S., Fitzkee N. C. Quantitative measurement of multiprotein nanoparticle interactions using NMR spectroscopy. *Anal. Chem.* **93**, 11982-11990 (2021).
10. Hubbard S., Thornton J. Naccess: Program for calculating accessibilities. *Department of Biochemistry and Molecular Biology, University College of London*, (1992).
11. Petkova G. A., Záruba K., Žvátora P., Král V. Gold and silver nanoparticles for biomolecule immobilization and enzymatic catalysis. *Nanoscale Res. Lett.* **7**, 287 (2012).
12. Korkhin Y., Kalb A. J., Peretz M., Bogin O., Burstein Y., Frolow F. Nadp-dependent bacterial alcohol dehydrogenases: Crystal structure, cofactor-binding and cofactor specificity of the adhs of clostridium beijerinckii and thermoanaerobacter brockii11edited by r. Huber. *J. Mol. Biol.* **278**, 967-981 (1998).
13. Baruah P., Yesylevskyy S. O., Aguan K., Mitra S. Modulation of enzyme activity at nano-bio interface: A case study with acetylcholinesterase and citrate synthase adsorbed on colloidal metal nanoparticles. *J. Mol. Liq.* **325**, 115201 (2021).
14. Yu X., Sigler S. C., Hossain D., Wierdl M., Gwaltney S. R., Potter P. M., et al. Global and local molecular dynamics of a bacterial carboxylesterase provide insight into its catalytic mechanism. *J. Mol. Model.* **18**, 2869-2883 (2012).
15. Remington S. J. Structure and mechanism of citrate synthase. In: Stadtman ER, Chock PB (eds). *Current topics in cellular regulation*, vol. 33. Academic Press, 1992, pp 209-229.
16. Cans A. S., Dean S. L., Reyes F. E., Keating C. D. Synthesis and characterization of enzyme-Au bioconjugates: HRP and fluorescein-labeled HRP. *NanoBiotechnology* **3**, 12-22 (2007).

17. Mogharrab N., Ghourchian H., Amininasab M. Structural stabilization and functional improvement of horseradish peroxidase upon modification of accessible lysines: Experiments and simulation. *Biophys. J.* **92**, 1192-1203 (2007).
18. Macdonald I. D. G., Smith W. E. Orientation of cytochrome c adsorbed on a citrate-reduced silver colloid surface. *Langmuir* **12**, 706-713 (1996).
19. Tellechea E., Wilson K. J., Bravo E., Hamad-Schifferli K. Engineering the interface between glucose oxidase and nanoparticles. *Langmuir* **28**, 5190-5200 (2012).
20. Leskovac V., Trivić S., Wohlfahrt G., Kandrač J., Peričin D. Glucose oxidase from *Aspergillus niger*: The mechanism of action with molecular oxygen, quinones, and one-electron acceptors. *Int. J. Biochem. Cell Biol.* **37**, 731-750 (2005).
